# Integrated spatial multi-omics reveals fibroblast fate during tissue repair

**DOI:** 10.1101/2021.04.02.437928

**Authors:** Deshka S Foster, Michael Januszyk, Malini S Chinta, Kathryn E Yost, Gunsagar S Gulati, Alan T Nguyen, Austin R Burcham, Ankit Salhotra, R Chase Ransom, Dominic Henn, Kellen Chen, Shamik Mascharak, Karen Tolentino, Ashley L Titan, R Ellen Jones, Oscar da Silva, W Tripp Leavitt, Clement D Marshall, Heather E desJardins-Park, Michael S Hu, Derrick C Wan, Gerlinde Wernig, Dhananjay Wagh, John Coller, Jeffrey A Norton, Geoffrey C Gurtner, Aaron M Newman, Howard Y Chang, Michael T Longaker

## Abstract

In the skin, tissue injury results in fibrosis in the form of scars composed of dense extracellular matrix deposited by fibroblasts. The therapeutic goal of regenerative wound healing has remained elusive in part because principles of fibroblast programming and adaptive response to injury remain incompletely understood. Here, we present a multimodal -omics platform for the comprehensive study of cell populations in complex tissue, which has allowed us to characterize the cells involved in wound healing across both time and space. We employ a stented wound model that recapitulates human tissue repair kinetics and multiple Rainbow transgenic lines to precisely track fibroblast fate during the physiologic response to injury. Through integrated analysis of single cell chromatin landscapes and gene expression states, coupled with spatial transcriptomic profiling, we are able to impute fibroblast epigenomes with temporospatial resolution. This has allowed us to define the mechanisms controlling cell fate during migration, proliferation, and differentiation following tissue injury and thereby reexamine the canonical phases of wound healing. These findings have broad implications for the study of tissue repair in complex organ systems.

---

Tissue fibrosis and its sequelae are associated with 45% of all mortality in the U.S^1, 2^. In the skin, wound healing is achieved through fibrosis and formation of a scar, which is composed of dense extracellular matrix components. Scars are stiff, poorly vascularized, and generally insensate. Scars also lack normal skin appendages (primarily hair follicles and sweat and sebaceous glands) and as such are devoid of the skin’s native functionality. As a result of these features, dermal scars can result in lifelong disability secondary to disfigurement and dysfunction^3^. Fibroblasts are responsible for the deposition of wound scar tissue. While several studies have characterized subtypes of fibroblasts involved in wound healing, the development of novel therapies that foster regeneration (rather than fibrosis) has remained limited because the origins, heterogeneity, and behavior of fibroblasts during tissue repair are not yet comprehensively understood.

Translating cutaneous tissue repair in mice to humans is challenging due to the presence of the panniculus carnosus muscle in mice. This is a subdermal muscle layer found throughout the body of the mouse that causes wound contraction, but whose analog in humans only exists as the platysma muscle in the neck, the palmaris brevis in the hand, and the dartos muscle in the scrotum. Fibroblast heterogeneity has been previously explored in wound healing using mouse models in which large, un-stented wounds (1.5 cm diameter) heal primarily by contraction with only a small portion in the center healing through re-epithelialization and deposition of connective tissue from fibroblasts (the primary mechanism of wound healing in humans)^4, 5^. To recapitulate clinically relevant wound healing using mouse models, we utilize a stented wound model which limits contraction of the panniculus carnosus and thereby mimics human wound healing kinetics^6^. Given that local tissue mechanics play a central role in scar formation^7–9^, this model permits us to interrogate fibroblast mechano-biology in a more clinically relevant manner.

Recent advances in sequencing and cell capture technology have enabled the assessment of gene expression with reference to tissue organization using spatial transcriptomics. This approach has only been applied to a limited number of tissue types to date, primarily in the study of tumors including prostate cancer^10^, skin cancer^11, 12^, and breast cancer^13^, as well as bone marrow^14^, joint inflammation^15^, and brain tissue^16^. However, to our knowledge, this technique has yet to be applied to characterize tissue repair or wound healing. Moreover, the spatial and temporal distributions of single cell chromatin landscapes – which mechanistically underlie gene expression – have yet to be described in any context.

Here, using transgenic mouse models, we assess the proliferation of local, tissue-resident fibroblast cells in wound healing. By establishing a microsurgical approach to independently isolate fibroblasts from spatially distinct regions within the wound, we interrogate Rainbow-labeled fibroblasts from critical timepoints during the course of wound closure. The Rainbow mouse model is a four-color reporter system that permits precise clonal analysis and lineage tracing. Using this model with phenotype-paired single cell RNA- and ATAC-sequencing (scRNA-seq and scATAC-seq), we are able to define the spatial and temporal heterogeneity of wound healing fibroblasts with unique granularity. Using full-length, plate-based scRNA-seq we assess the differentiation states of individual cells as they proliferate and migrate from the outer wound region inward^17^. By disrupting this process using small molecule inhibition or genetic knockdown of focal adhesion kinase (FAK, *Ptk2*), we further elucidate the relationship between wound healing fibroblast activation and microenvironmental cues. By integrating our scRNA-seq and scATAC-seq analyses using the recently-developed ArchR platform^18^, we delineate interrelated changes in chromatin accessibility and gene expression driving wound closure and scar fibrosis and identify functionally distinct wound healing fibroblast subpopulations. Furthermore, using CIBERSORTx deconvolution^19^ of bulk RNA-seq data, we are able to categorize putative fibroblast subpopulations-based response to local tissue injury. Finally, we introduce spatial multi-omics, combining spatial transcriptomics with paired scRNA-seq and ATAC-seq datasets to impute spatial epigenomic properties. These data provide a comprehensive map of fate-determining chromatin accessibility states in the healing wound. Collectively, this work defines the spatial and temporal dynamics of the cellular response to injury and provides a multimodal -omics framework for future studies in tissue repair.

## Wounding triggers polyclonal proliferation of tissue-resident fibroblasts

Many cell surface and lineage markers have been associated with fibroblasts involved in wound healing, including *Pdgfra*, Engrailed-1 (*En1*), and CD26 (*Dpp4*)^20, 21^. However, we and others have found expression of such markers to be highly variable throughout wound tissue (**Extended Data Fig. 1a**), suggesting spatial and functional heterogeneity among the fibroblasts that respond to injury. We asked whether there might be one or more fibroblasts activated following injury that could give rise to more diverse downstream fibroblast phenotypes. If so, we wondered whether such a cell type would be of tissue-resident origin, as suggested by previous studies^22–24^ (**Extended Data Fig. 1b**), or originate from the peripheral circulation. To explore this, we employed transgenic parabiotic mice in conjunction with an established model of excisional wound healing^6^ (**Extended Data Fig. 1c**). Each eGFP donor mouse was parabiosed to a wild-type (C57BL/6J) mouse (**Extended Data Fig. 1d**), and shared blood supply was established by two weeks after surgery (**Extended Data Fig. 1e**). At that time, wounds were made on the dorsum of each wild-type parabiont and then harvested at either post-operative day (POD) 7 (midway through healing) or POD 14 (when the wound has fully re-epithelialized). While systemically infiltrating GFP+ cells were found in the dermal scar at both time points, the overwhelming majority of GFP+ cells were also CD45+ and thus of hematopoietic (non-fibroblast) lineage (**Extended Data Fig. 1f**). These data further support the growing body of literature indicating that the fibroblasts responsible for wound healing are local, tissue-resident cells.

**Figure 1.**
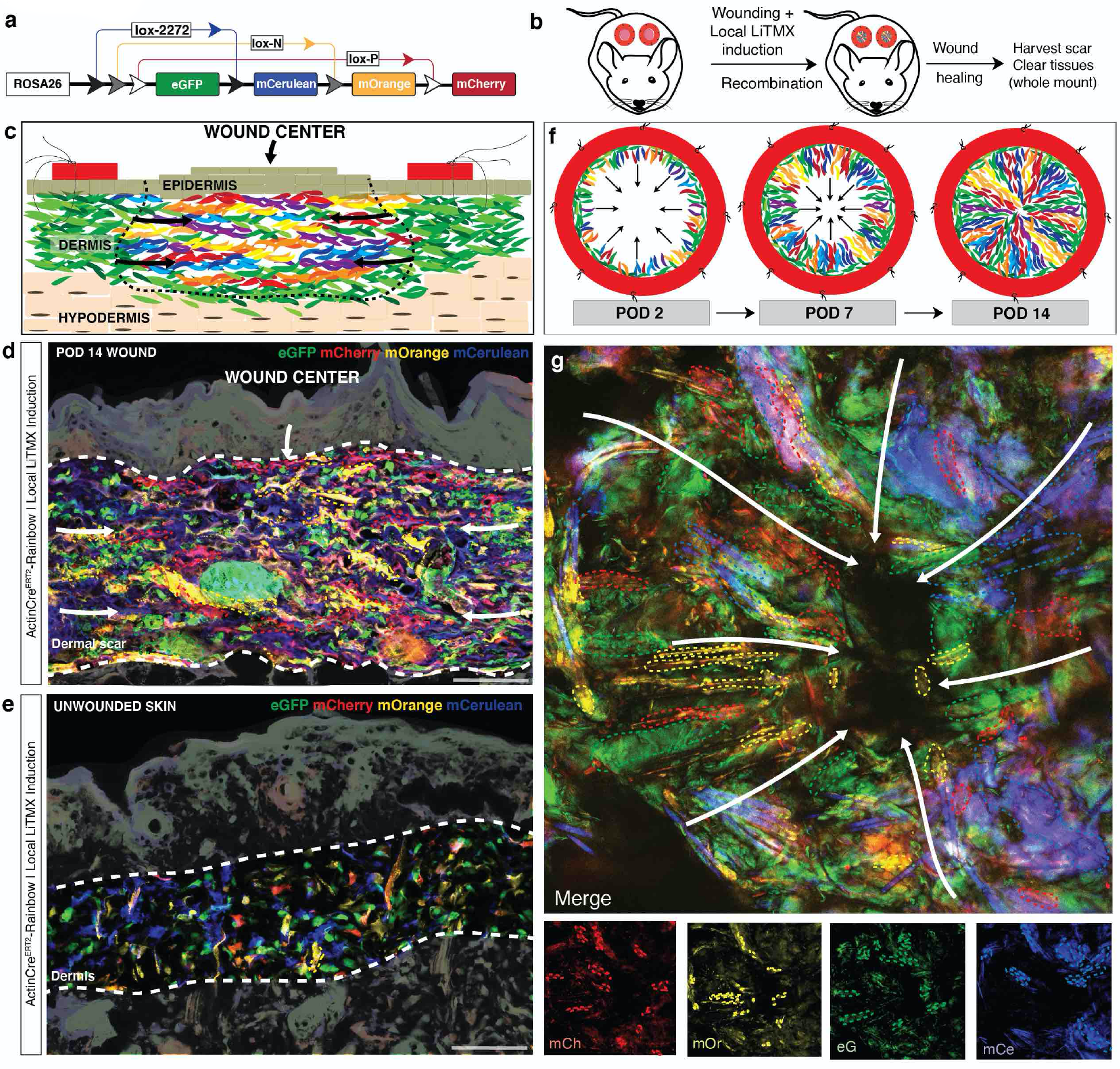
Wounding triggers polyclonal proliferation of tissue-resident fibroblasts. **a**, Schematic of the Rainbow mouse construct. **b,** Schematic showing wound healing model using Rainbow mice with local Cre recombinase induction using 4-hydroxytamoxifen liposomes (LiTMX). **c,** Schematic showing a Rainbow wound cross-section. Black dotted line highlights wound scar area; arrows indicate the direction of cellular proliferation during wound healing. Structures as labelled in panel. **d,** Representative confocal image of POD 14 wound cross sections from *Actin-Cre^ERT^*^2^*::Rosa26^VT^*^2^*^/GK^*^3^ mice induced locally with LiTMX at the time of wound creation. Thick white dotted lines highlight scar boundaries. Individual Rainbow cell clones are highlighted with thin colored dotted lines. Arrows indicate direction of wound healing. *n > 5.* Scale bar = 50μm. **e,** Representative confocal images of unwounded skin from *Actin-Cre^ERT^*^2^*::Rosa26^VT2/GK3^* mice induced locally with LiTMX. Thick white dotted lines highlight dermal boundaries. Individual Rainbow cell clones are highlighted with thin colored dotted lines. *n > 5.* Scale bar = 50μm. **f,** Schematic of dorsal, stented, excisional wound healing in the Rainbow mouse model (whole mount view), with polyclonal proliferation of Rainbow fibroblasts from the outer wound edge inward across time from POD 2 (**left panel**), to POD 7 (**middle panel**), to POD 14 (**right panel**). Black arrows highlight the apparent direction of proliferation. **g,** Representative confocal imaging of a POD 14 whole-mounted wound harvested from *Actin-Cre^ERT2^::Rosa26^VT2/GK3^* mice showing the polyclonal proliferation of wound fibroblasts radially towards the center of the wound (dark area at center). White arrows highlight the direction of cell proliferation; individual cell clones are highlighted with thin colored dotted lines. Bottom subpanels denote individual Rainbow color contributions to merged image. mCh = membrane (m)Cherry, mOr = mOrange, mCe = mCerulean, eG = eGFP. *n* > 5.

To further explore the lineage dynamics, activation, and proliferation of these cells, we examined stented wound healing using the Rainbow (*Rosa26^VT^*^2^*^/GK^*^3^) mouse model^25^. Rainbow mice contain a transgenic four-color reporter construct in the *Rosa26* locus. Upon induction with Cre-recombinase, the four colors irreversibly recombine (creating up to ten color combinations) and all subsequent progeny cells will have the same color as their parent cells, permitting stochastic lineage tracing and clonal analysis (**Fig. 1a**). We developed a technique using local induction with activated tamoxifen liposomes (LiTMX) to exclusively consider tissue-resident cells (**Fig. 1b**)^26^. Following injury, tissue-resident fibroblasts were found to proliferate in a linear, polyclonal manner along the cross-sectional wound interface (**Fig. 1c-d**), whereas uninjured dermal fibroblasts exhibited only minimal clonality (**Fig. 1e**). These data support the presence of local, cells that are activated in response to injury and proliferate polyclonally to fill the wound “gap.” We developed an optimized tissue clearing protocol and whole mount technique to comprehensively visualize wound healing biology with the Rainbow mouse^27^. Using these methods in conjunction with a ubiquitous *Actin-Cre^ERT^*^2^ driver, we observed that cells were activated along the wound edge and proliferated inward in a distinct radial pattern (**Fig. 1f-g**).

## Bulk transcriptomic analysis of injury-responsive fibroblasts

Based on the pattern of clonal proliferation extending from the outer wound edge inward, we sought to determine whether there might be underlying, region-specific cellular changes that are characteristic to this process. As such, we developed a microsurgical isolation technique that considers the “inner” and “outer” components of the wound dermis independently (**Fig. 2a**). We isolated wound fibroblasts from these two regions separately at POD 7 (midpoint of healing). Bulk RNA-seq evaluation showed clear differences in the gene expression profiles of both inner and outer wound fibroblasts compared with those of uninjured (control) skin (**Extended Data Fig. 2a**). We noted upregulation of fibrosis genes such as matrillin (*Matn2*), *Dlk1*, osteopontin (*Spp1*), *Acta2,* and multiple collagen subtypes (**Extended Data Fig. 2b-c**). We also observed elevated expression of mechanotransduction and FAK pathway components such as the chemokines *Stat1* and *Il6*, consistent with our prior work demonstrating FAK regulation of cell-matrix interactions in wound healing^28^. When we directly compared inner and outer wound fibroblasts (**Fig. 2b**), we found that cell cycle pathways were among those upregulated in inner compared to outer region wound fibroblasts (**Fig. 2c** **– top panel**), whereas outer fibroblasts had greater enrichment for more generalized tissue-repair processes (**Fig. 2c** **– bottom panel**). Furthermore, we observed that inner wound fibroblasts had transcriptional programs more divergent from uninjured skin than their outer wound counterparts (**Extended Data Fig. 2c**). These findings support broad regional differences in the proliferation and activation status of fibroblasts in the healing wound; however, their strength is limited by the lack of granularity inherent to bulk transcriptional analysis.

**Figure 2.**
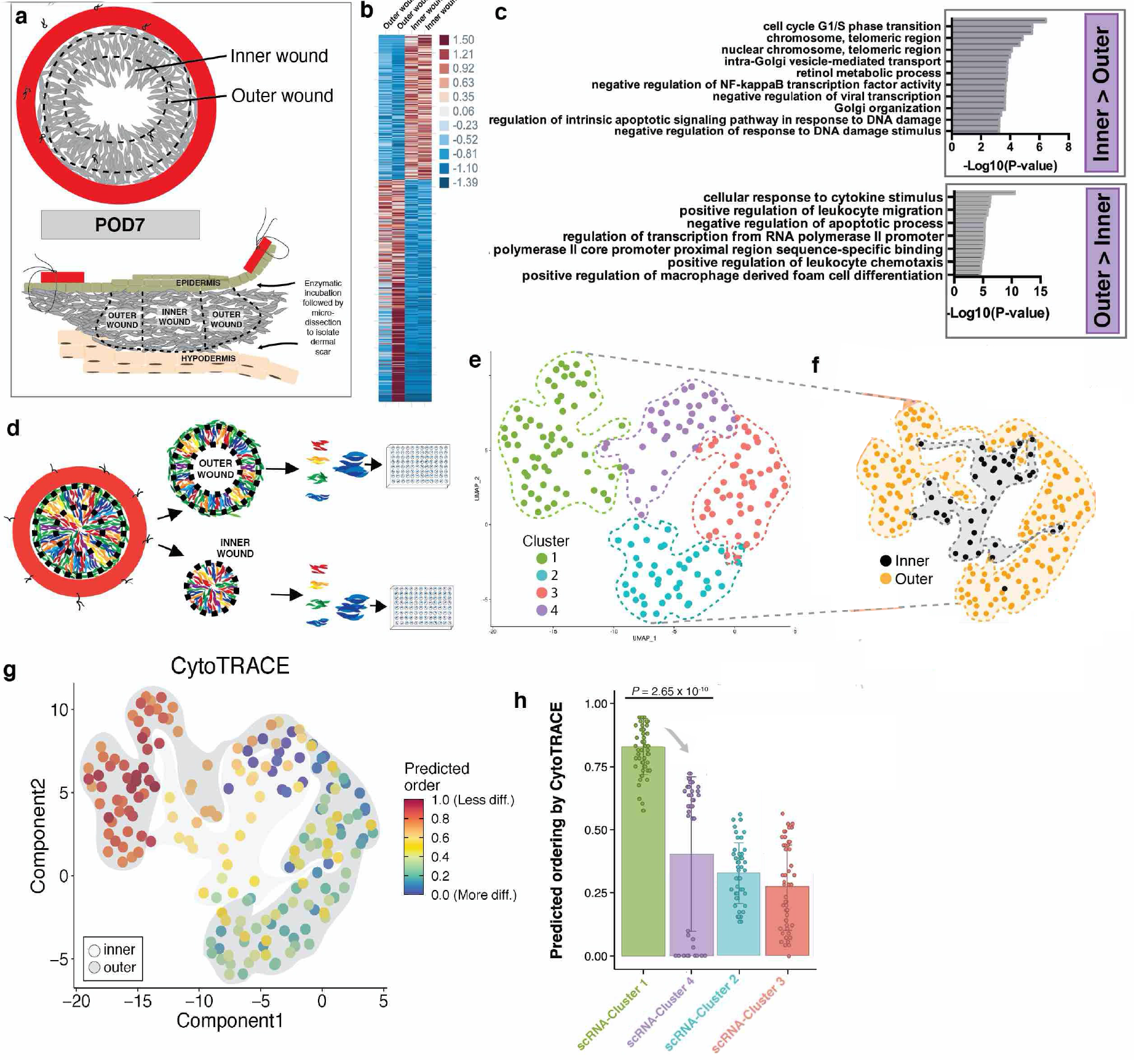
Bulk and single-cell transcriptomic analysis of injury-responsive fibroblasts. **a,** Schematics illustrating micro-dissection strategy for isolation of inner and outer wound regions (**top panel**), followed by enzymatic separation of the dermal scar from the epi- and hypo-dermis (**bottom panel**). **b,** Heatmap displaying expression data for genes significantly different between POD 7 inner and outer region wound fibroblasts. Legend at right displays fold-change. **c,** Gene Ontology (GO) enrichment analysis comparing gene expression data from POD 7 inner and outer region wound fibroblasts. Top panel shows GO Biological Processes upregulated in inner region fibroblasts compared with outer region fibroblasts, while the bottom panel shows the same for outer region fibroblasts compared with inner. Top 10 most significant gene sets are displayed for each condition. **d,** Schematic illustrating single-cell (sc) isolation of Rainbow wound fibroblasts from inner and outer wound regions (highlighted with black dotted lines). For scRNA-seq, mCerulean+ fibroblasts were arbitrarily selected from the available Rainbow colors and used for the remaining experiments in this Figure. **e,** Uniform manifold approximation and projection (UMAP) embedding showing scRNA -seq data from mouse wound fibroblasts FACS-isolated using a lineage-negative sort strategy ^29^ from POD 2, POD 7, and POD 14 - digitally pooled and clustered in a manner agnostic to POD and inner vs outer wound regions. Four unique fibroblasts clusters were identified (Cluster 1-4). Dotted lines highlight individual clusters distributions. **f,** Re-coloring of Fig. 2e UMAP plot based on fibroblast tissue region: inner (black) versus outer (orange). **g,** CytoTRACE analysis of scRNA-seq data using the UMAP embedding from Fig. 2f. Shading indicates inner (light grey) versus outer (dark grey) wound regions. **h,** Box plots showing the predicted ordering by CytoTRACE for individual cells within the four scRNA-seq clusters. Grey arrow indicates direction of predicted differentiation from scRNA-seq Cluster 1 to Cluster 4 (which corresponds to outer-to-inner wound region expansion). *P*-value was derived from two-sided Student’s *t*-test.

## Traditional cell surface markers are not sufficient to characterize regional heterogeneity among wound healing fibroblasts

We evaluated how well several recently published cell surface marker profiles, defining fibroblast subtypes largely based on tissue depth, tracked with the regional differences observed in our study^20^. FACS-isolated, lineage-negative ^29^, Rainbow wound fibroblasts were examined (**Fig. 2d, Extended Data Fig. 3a**), and we found that most of cells fell into the putative category of reticular fibroblasts (defined as DLK1^+^/SCA1^-^) rather than papillary (CD26^+^/SCA1^-^) or hypodermal (DLK1^+/-^/SCA1^+^) (**Extended Data Fig. 3b**). When we considered inner and outer region wound fibroblasts separately, we found that distribution of fibroblast subtypes was not significantly different between these two groups (**Extended Data Fig. 3c**), suggesting that differences in expression among known marker profiles are not sufficient to delineate inner versus outer region wound fibroblasts, though these can be readily distinguished based on their transcriptional programs even at the bulk tissue level.

**Figure 3.**
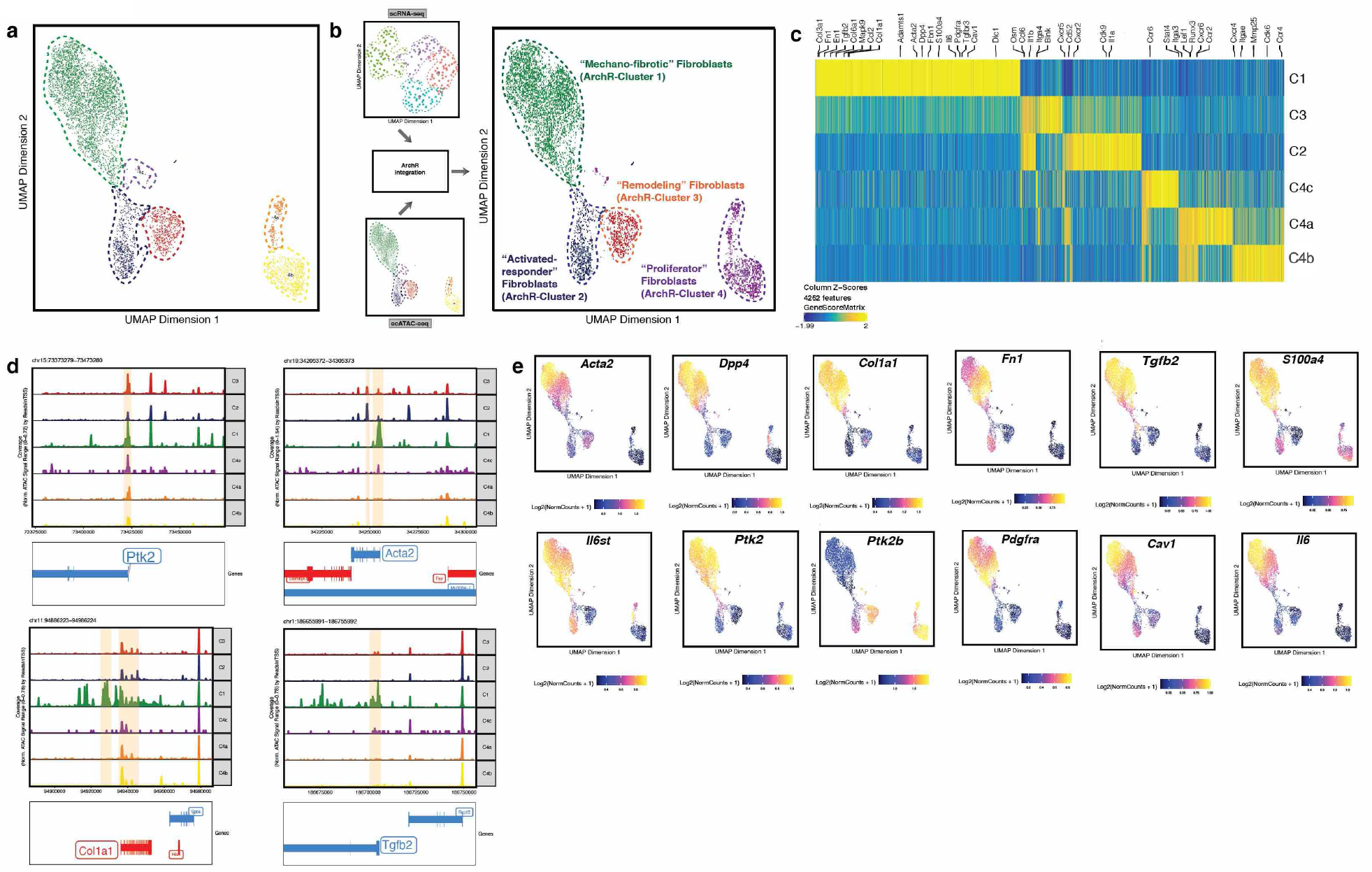
Chromatin accessibility delineates mechano-responsive fibroblast subpopulations. **a,** scATAC-seq evaluation of Rainbow mouse wound fibroblasts isolated in parallel with our scRNA-seq experiments (see Methods). UMAP embeddings were generated using ArchR with default Louvain parameters^18^, and six unique fibroblast clusters were identified. **b,** ArchR integration of scRNA-seq and scATAC-seq data. **c,** Heatmap of scATAC-seq cluster peaks mapped to associated gene markers. Elements relevant to fibroblast activation, fibrosis, and mechanotransduction are specifically annotated along the top of panel. **d,** Genome tracking plots showing scATAC-seq peaks for pseudo-bulk replicates generated for each cluster. Associations between the peaks with fibrosis and mechanotransduction-related genes (Peak2GeneLinks) are included at the bottom of each plot. Pale orange shading highlights differentially expressed peaks across the scATAC Clusters. All highlighted peaks demonstrated statistically significant differential expression in at least one pairwise comparison (FDR < 0.1 and FC >= 2). **e,** UMAP feature plots highlighting distributions for genes of interest related to fibrosis and mechanotransduction.

## Single-cell transcriptomic analysis of injury-responsive fibroblasts

We sought to better characterize wound fibroblast heterogeneity by examining individual fibroblast transcriptional programs at important functional timepoints in the canonical wound healing process: POD 2 – inflammation, POD 7 – granulation, and POD 14 – complete re-epithelialization (“healed” wound). We conducted single-cell, plate-based RNA-seq (scRNA-seq) of lineage-negative fibroblasts isolated based on their expression of Rainbow clone colors from both inner and outer wound regions at each time point (**Fig. 2d**). Modularity-optimized clustering of the resulting gene expression dataset identified four transcriptionally-defined fibroblast subpopulations (**Fig. 2e**), with considerable differences in their distributions between wound regions (**Fig. 2f**).

Given our interest in understanding lineage trajectories in the context of wound healing, we assessed the relative differentiation states of these fibroblast populations using CytoTRACE, a computational tool that leverages transcriptional diversity to order cells based on developmental potential (**Fig. 2g, Extended Data Fig. 4**)^17^. This analysis identified a clear lineage trajectory stemming from scRNA-Cluster 1, which is primarily represented by cells from the outer wound region, extending to scRNA-Cluster 4, which is primarily represented by cells from the inner wound (**Fig. 2h**). These findings suggest that fibroblasts undergo differentiation and activate fibrotic transcriptional programs as they proliferate from the outer wound inward.

**Figure 4.**
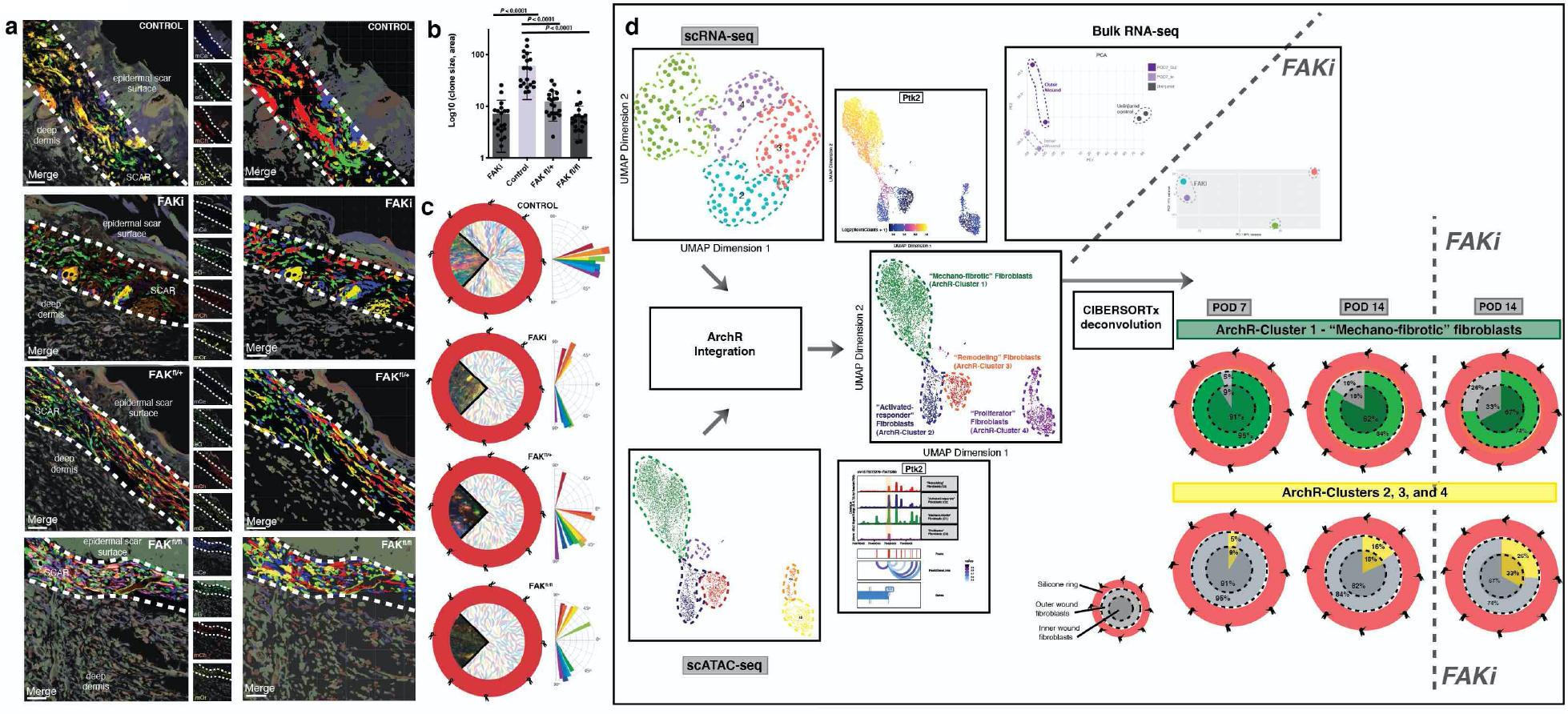
Clonal proliferation of injury-responsive fibroblasts is dependent on mechanotransduction signaling. **a,** Representative confocal images of sectioned Rainbow mouse wound specimens treated with FAKi (**second panels**), FAK^fl/+^ (**third panels**) or FAK^fl/fl^ (**bottom panels**) compared with vehicle control (**top panels**). Imaris rendering in second column of images highlights individual Rainbow clones. Dermal wound area highlighted with thick while dotted line. *n* = 5. Scale bars = 25μm. **b,** Quantitation of average clone size based on Imaris rendering. **c,** Wedge sections of representative whole mount confocal images of Rainbow wound specimens embedded within surrounding wound schematics for vehicle control (**top panel**), FAKi-treated (**second panel**), FAK^fl/+^ (**third panel**), and FAK^fl/fl^ (**bottom panel**) samples. Corresponding vector analyses are provided to the right of each sub-panel. **d,** Schematic illustrating our approach to deconvolve bulk RNA-seq data using our multimodal scRNA-ATAC construct. Transcriptionally-defined cluster labels from scRNA-seq analysis were projected onto the scATAC-seq manifold using an anchor transfer-based approach in ArchR as previously described^18^ to construct four multimodal fibroblast subpopulations (**left-center panel**). Putative names were assigned to these ArchR-Clusters based on integrated functional and temporospatial characteristics. Feature and peak plots, above and below, for FAK (*Ptk2)* are provided for illustrative purposes. Deconvolution of bulk RNA-seq specimens representing wound fibroblasts treated with FAKi versus vehicle control (**middle panel**) was then performed using CIBRERSORTx^19^ (see Methods). Wound schematics (with silicone ring around the outside, and outer and inner regions indicated) are provided to represent CIBRERSORTx output identifying changes in the percentages of ArchR-Cluster 1 (“Mechano-fibrotic”) cells in bulk samples over time and with/without FAKi treatment (**green, upper row of plots**). Parallel schematic of corresponding changes in other ArchR-Clusters are provided in yellow (**lower row of plots**).

## Evaluation of chromatin accessibility complements transcriptional analysis of mechano-responsive fibroblast subpopulations

To evaluate the epigenomic changes associated with fibroblast activation and lineage differentiation in wound healing, we conducted a series of scATAC-seq experiments in parallel with our scRNA-seq assays (**Extended Data Fig. 5a**). We identified considerable heterogeneity in accessibility profiles among individual wound fibroblasts, which were clustered into six epigenomically-distinct subgroups using a Louvain-based modularity optimization in the ArchR platform^18^ (**Fig. 3a**). This partitioning was agnostic to the phenotype of cell origin (i.e., wound region or post-operative day), and all clusters included fibroblasts harvested from multiple time points and wound regions (**Extended Data Fig. 5b-c**).

**Figure 5.**
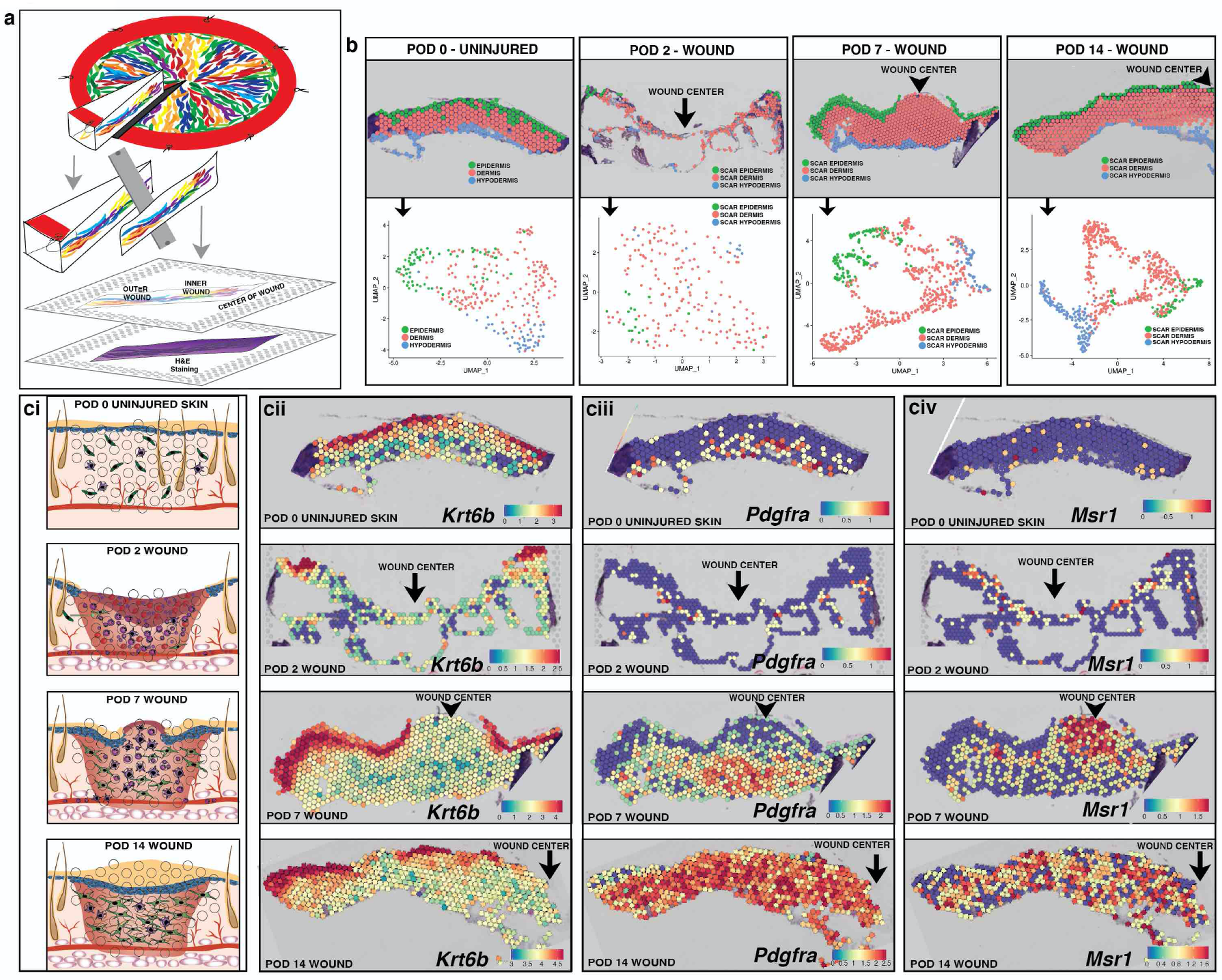
Spatial transcriptomics applied to wound healing. **a,** Schematic for generating spatial transcriptomics data from splinted excisional wounds using the 10x Genomics Visium protocol. Fresh Rainbow mouse wound tissue is harvested, flash-frozen embedded in OCT, and then sections were taking representing the complete wound radius. H&E staining and tissue section imaging was completed as described in the Visium protocol (see Methods). Each “spot” captures mRNA from 1-10 individual cells at that tissue location. **b,** Delineation of scar layers based on underlying tissue histology at each time point (top row), and UMAP plot showing that the three scar layers can easily be distinguished by their transcriptional programs, even independent of spatial information. **c,** (**i**) Schematic of classic stages of wound healing evaluated at POD 2, 7, and 14 relative to uninjured skin. (**ii**) Keratinocyte activity as measured through expression of the *Krt6b* gene. (**iii**) Fibroblast activity as measured through expression of the *Pdgfra* gene. (**iv**) Immune cell activity as measured through expression of the *Msr1* gene.

We then performed cross-platform integration to link these scATAC data with our earlier scRNA data using ArchR’s phenotype-constrained implementation of Seurat’s label transfer algorithm^18^. This resulted in four multimodal clusters characterized by both gene expression and chromatin accessibility profiles (**Fig. 3b, Extended Data Fig. 5d**), which we refer to as ArchR-Clusters 1-4 (**Extended Data Fig. 6a-b**).

**Figure 6.**
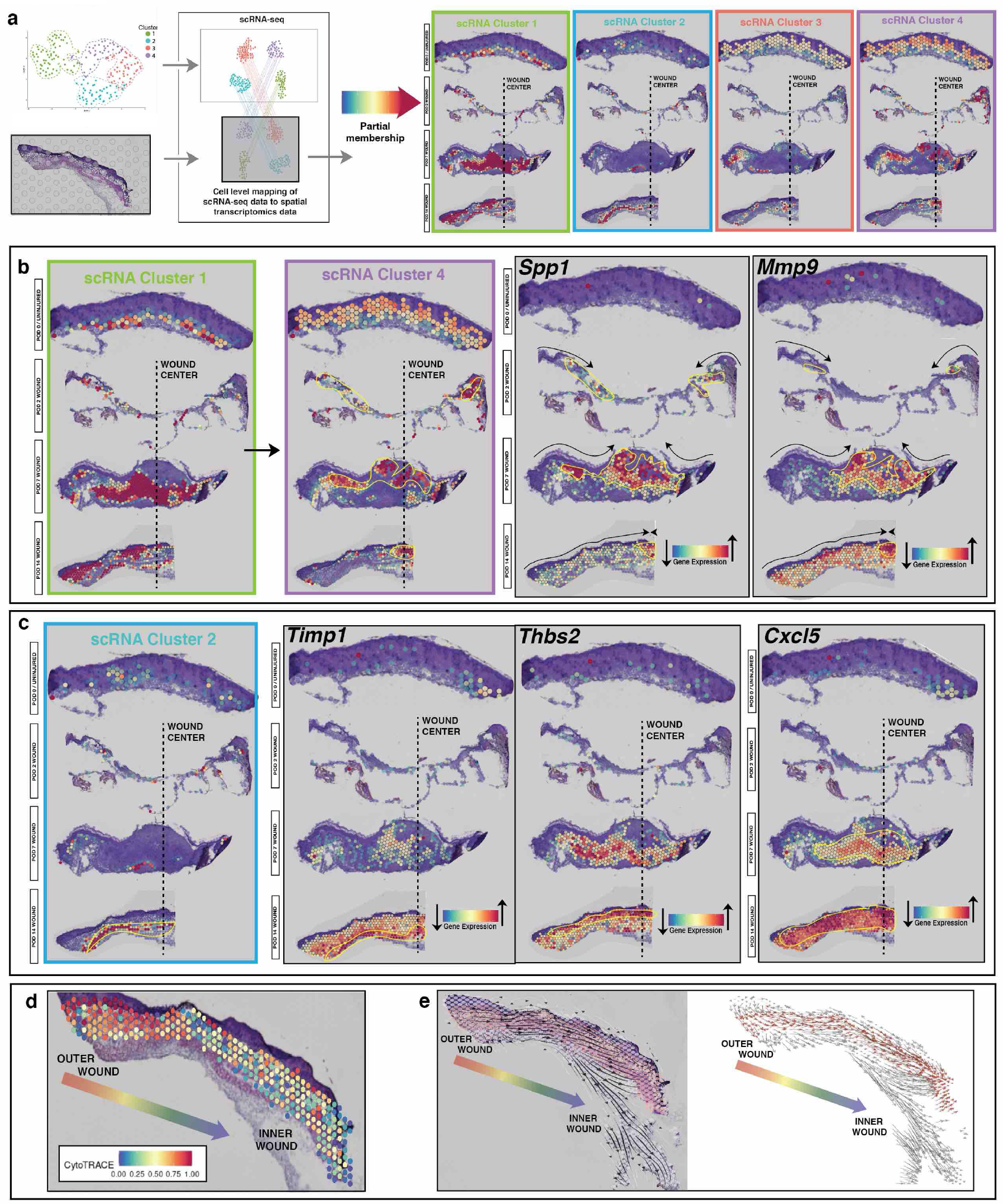
Tracking fibroblast subpopulations over time and space. **a,** Anchor-based integration of scRNA-seq populations (defined in Fig. 2f) with Visium gene expression to project partial membership within each spot across all time points. These populations exhibit strong spatial preferences within the wound. **b,** Visium plots showing POD 0, 2, 7, and 14 (top to bottom) wound sections. As wound fibroblasts proliferate from the outer region of the wound inwards, a portion of Cluster 1 cells appear to differentiate into Cluster 4 cells. Cluster 4 cells prominently express Spp1 and Mmp9, and track along the same trajectory of the healing edge of the epidermis. Wound centers indicated by dotted line as labelled in panels. Left 2 panels show partial membership clusters from a. Right two panels show SCT gene expression. **c,** Visium plots showing POD 0, 2, 7, and 14 (top to bottom) wound sections. Cluster 2 cells appear to populate the basal dermis primarily. Gene expression of Timp1 highlights this subpopulation, whereas Thbs1 expression highlights suprabasal wound fibroblasts, which strongly express chemokines such as Cxcl5 regulating immune cells within the wound microenvironment. Wound centers indicated by dotted line as labelled in panels. Left panel show partial membership clusters from a. Right three panels show gene expression data. **d,** CytoTRACE applied to spatial transcriptomics recapitulates cellular differentiation from the outer to inner regions of the wound. **e,** RNA-velocity data demonstrating a clear arc of differentiation from outer to inner wound regions along the dermal scar layer.

We first examined the epigenomic landscape of the largest subpopulation, ArchR-Cluster 1, which showed significantly elevated chromatin accessibility proximal to key fibrosis-related genes such as *Col1a1, Acta2,* and *Pdgfra* (**Fig. 3c-e, Extended Data Fig. 6c-e**). We also observed specific accessibility peaks and transcription factor footprinting in association with the FAK (*Ptk2*) locus, as well as numerous FAK-pathway elements, suggesting that these fibroblasts represent a mechanoresponsive, pro-fibrotic subpopulation (**Extended Data Fig. 7a-b, Extended Data Fig. 8**). ArchR-Cluster 2 was associated with elevated *Fn1* and *Thbs1* accessibility peaks; ArchR-Cluster 3 was characterized by increased accessibility at the *Jak2* locus and decreased accessibility at the *Fsp1 (S100a4*) and *Il6st* loci; and ArchR-Cluster 4 was characterized by increased accessibility at the *Ptk2b, Jak1*, and *Jak3* loci.

**Figure 7.**
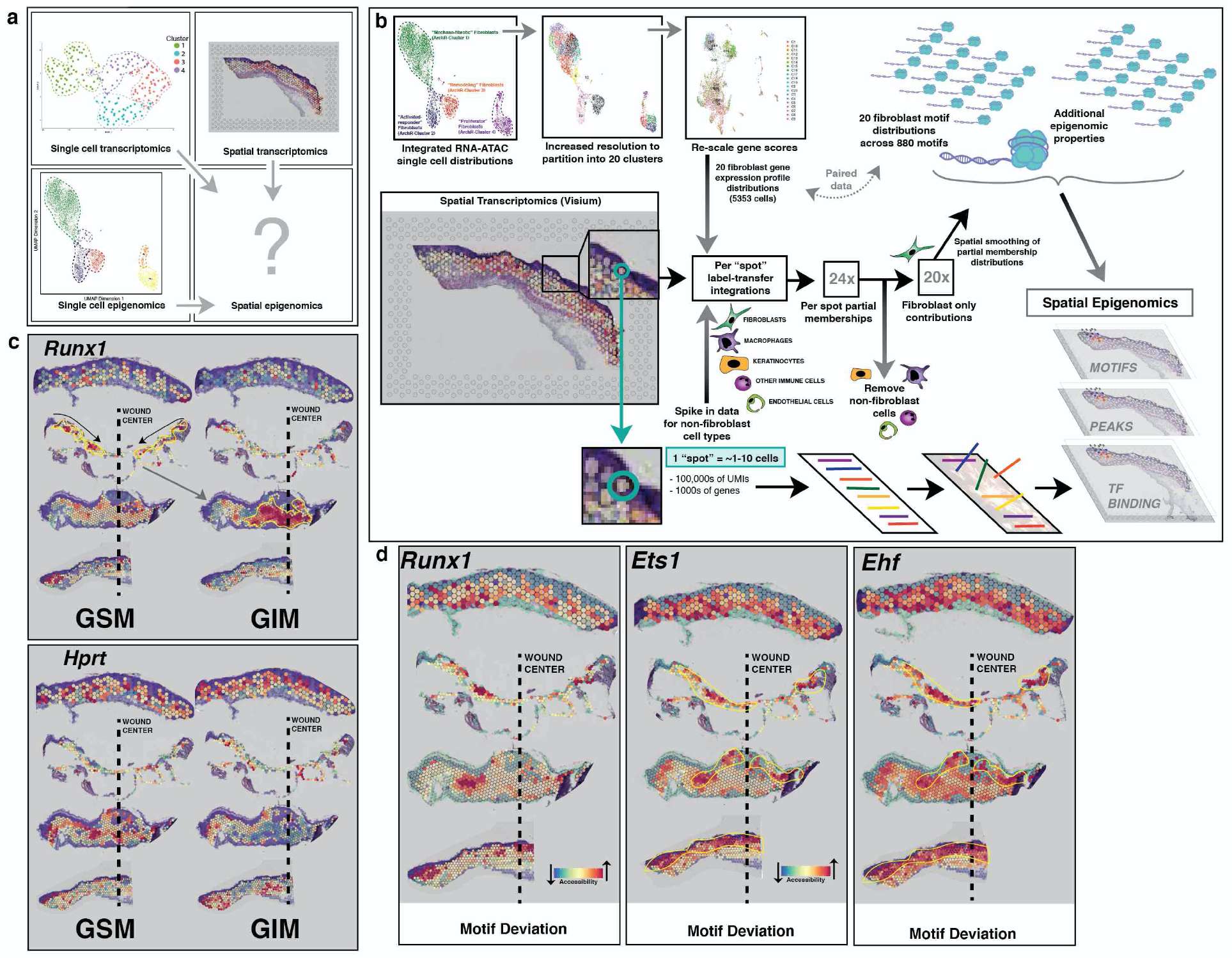
Integrated analysis permits imputation of spatial epigenomic properties. **a,** Punnett square schematic summarizing the data acquired in Figures 2, 3, and 6; setting the stage for imputation of spatial epigenomics. **b,** Schematic summarizing imputation of spatial epigenomics. Multimodal scRNA-ATAC fibroblast data were first re-clustered into a higher-resolution space to generate twenty partitions, each representing between 27 and 552 cell equivalents. Gene score matrix distributions, informed by both modalities, were then extracted for each partition and subjected to SCT transformation. “Spike-in” RNA-seq data for keratinocytes, endothelial cells, granulocytes, and macrophages were obtained pure Visium spots across all time points. These data were combined and subjected to a similar variance-stabilizing transformation. The resulting putative single cell gene expression reference matrix was then used to assign initial partial set memberships for each spatial transcriptomic datapoint using an anchor transfer-based approach. Non-fibroblast contributions were subsequently regressed out, and a single-step spatial smoothing filter was applied to the resulting membership space, followed by re-normalization. The resulting partial set memberships for each spatial datapoint were then treated as a topological vector space, onto which epigenomic peak, motif, and binding activity from the twenty scRNA-ATAC partitions can be projected. **c,** Visium plots showing POD 0, 2, 7, and 14 (top to bottom) wound sections, imputed spatial epigenomics. For housekeeping genes such as *Hprt* (top panel), gene imputed matrix (GIM) correlates with gene score matrix (GSM) epigenomic data and is fairly stable over space and time (**top panel)**. However, for *Runx1*, which we have shown to be very active within wound fibroblasts, GSM data shows opening at the *Runx1* motif at POD2, which yields strong gene expression primarily among inner wound fibroblasts at POD 7 (**bottom panel**). **d,** Visium plots showing POD 0, 2, 7, and 14 (top to bottom) wound sections, motif deviations for genes of interest related to FAK-mediated mechanotransduction and fibroblast proliferation including *Runx1, Ets1,* and *Ehf*.

In addition to specific peak and motif evaluation, we also employed cluster-wide enrichment analysis using Genomics Regions Enrichment of Annotations Tool (GREAT)^30^ (**Extended Data Fig. 9a**). We found significant enrichment for “Focal adhesion” in ArchR-Cluster 1 and for FAK-pathway signaling response elements such as “Integrin a11b1” in ArchR-Clusters 1 and 4. The related processes of “Increased fibroblast migration” and “Increased fibroblast proliferation” were specifically enriched for among Cluster 1 and Cluster 4 cells, respectively. Furthermore, pseudotime analysis of these integrated scRNA-ATAC data demonstrated an epigenomic progression from the putatively least-differentiated ArchR-Cluster 1 to the remaining cell populations that was driven by mechanical signaling elements (**Extended Data Fig. 9b**).

Based on these findings, we were able to provisionally characterize each subpopulation according to its putative role in the wound healing process: “Mechano-fibrotic” (ArchR-Cluster 1), “Activated-responder” (ArchR-Cluster 2), “Remodeling” (ArchR-Cluster 3), and “Proliferator” (ArchR-Cluster 4) fibroblasts (**Fig. 3b, right**).

## Clonal proliferation of injury-responsive fibroblasts is mechanotransduction-dependent

Our laboratory has previously shown that local tissue mechanics are crucial in guiding the response to healing after injury^31^, and mechanotransduction signaling pathway elements were found to delineate fibroblast subpopulations in our scRNA and scATAC wound data. To further interrogate the role of local tissue mechanics in wound biology, we applied a small molecule FAK inhibitor (FAKi) to disrupt mechanosensation in stented mouse wounds (**Extended Data Fig. 10a**). Consistent with prior work, we found that FAKi-treated wounds healed at the same rate as untreated wounds (**Extended Data Fig. 10b-c**), but resulted in significantly smaller and thinner scars composed of less-dense matrix tissue (**Extended Data Fig. 10d**) and connective tissue arranged in a more basket-weave-like pattern characteristic of uninjured skin (**Extended Data Fig. 10e**)^28^.

To validate our FAKi results, we conducted additional wound healing experiments using *Actin-Cre^ERT^*^2^*::Rosa26^VT^*^2^*^/GK^*^3^*::Ptk2^fl/+^* and *Actin-Cre^ERT^*^2^*::Rosa26^VT^*^2^*^/GK^*^3^*::Ptk2^fl/fl^* (heterozygous [*FAK^fl/+^*] and homozygous [*FAK^fl/fl^*] knock-out) mice, with local LiTMX induction at the time of wounding (**Extended Data Fig. 10a-c**). We found that these mouse wounds also exhibited a less scar-like pattern of connective tissue (**Extended Data Fig. 10e**). To further explore these differences, we employed an automated feature extraction algorithm (Mascharak et al., *in review*) to identify and quantify hundreds of ultrastructure characteristics for Picrosirius red stained wound tissue sections. Lower dimensional embedding of these features demonstrated that FAKi-treated or knock-out wound specimens were more similar to unwounded skin than to vehicle-control wounds, with significant differences present for both mature and immature collagen fiber intensities (**Extended Data Fig. 10f**). Taken together, these findings confirm that when FAK is inhibited, either genetically or using a small molecule, wounds heal with thinner scars and connective tissue structure that is more similar to that of unwounded skin.

To understand the transcriptional changes associated with modulation of mechanotransduction in wound healing, we conducted additional bulk RNA-seq experiments comparing fibroblasts isolated from inner and outer regions of FAKi-treated and control wounds. We observed significant changes in the transcriptional programs of FAKi-treated cells, and found that the regional differences were diminished in wounds following FAK inhibition (**Extended Data Fig. 11a-b**). These results suggest that local tissue mechanics play a key role in the underlying changes to transcriptional programming between inner and outer wound regions. We also found that wound healing fibroblasts showed downregulation of mechanotransduction and fibrosis-related pathways with FAKi treatment **(Extended Data Fig. 11c**), again supporting the conclusion that disruption of FAK signaling impedes fibroblast scarring capacity.

We next applied FAKi to Rainbow mouse wounds to explore the effect of FAK modulation on the polyclonal proliferation of fibroblasts. We found that when mechano-signaling was blocked using FAKi, or in *FAK^fl/+^* or *FAK^fl/fl^* mice, the linear polyclonal proliferation of fibroblasts that was previously appreciated (**Fig. 1g**) was disrupted (**Fig. 4a**). With FAKi, or in *FAK^fl/+^* or *FAK^fl/fl^* mice, the resulting rainbow fibroblast clones were smaller and less ordered (**Fig. 4b-c**).

## Deconvolution permits evaluation of bulk tissue samples through the lens of single cell -omics

We applied the deconvolution tool CIBERSORTx^19^ to estimate the abundance of our four scRNA-ATAC populations (ArchR-Clusters 1-4) within bulk RNA-seq data for fibroblasts isolated from POD 7 and POD 14 wounds with or without FAKi treatment (**Fig. 4d**). We found that the majority of cell estimates across the specimens were attributed to “Mechano-fibrotic” ArchR-Cluster 1, consistent with its prominent representation in both our scRNA-seq and scATAC-seq datasets, likely comprising among the most active fibroblasts during wound healing. The predicted prevalence of these cells was highest at POD 7 (91% and 95% in the inner and outer regions, respectively) and decreased by POD 14 (82% and 84%). FAK inhibition demonstrated a more dramatic reduction of ArchR-Cluster 1 fibroblasts at POD 14 for both inner (67%) and outer (74%) wound samples, further supporting the mechano-sensitivity of the ArchR-Cluster 1 fibroblast subpopulation.

## Spatial transcriptomics applied to wound healing

To further explore the significance of fibroblast heterogeneity in healing wounds, we applied the recently developed 10x Genomics Visium platform to analyze gene expression while retaining tissue spatial information. This technology had not previously been applied to analyze skin wounds, so we first developed an optimized protocol that permitted us to obtain reproducible, high-quality spatial gene expression data for all stages in the healing process (see Methods). We then conducted spatial transcriptomic analysis on tissue from our stented Rainbow mouse wound healing model at POD 2, 7, and 14, as well as in uninjured skin (**Fig. 5a**).

The epidermal, dermal, and hypodermal layers of the healing wounds were easily delineated histologically and also found to cluster independently based on transcriptional programs (**Fig. 5b**). Looking at individual genes for prominent wound healing cell types (**Fig. 5ci**), we found clear delineation of keratinocytes in the epidermis based on *Krt6b* expression (as well as other keratinocyte-specific gens), allowing us to track re-epithelialization over space and time at the transcriptional level (**Fig. 5cii**). Similarly, fibroblast activity was evaluated using characteristic genes such as *Pdgfra*, which was most prominent in the dermis and most active at POD 14 (**Fig. 5ciii**). Likewise, we found that activated macrophage markers like *Msr1* allowed us to monitor these immune cells throughout our dataset, which were very prominent in the “proud flesh” at the center of the wound at POD 7 (**Fig. 5civ**).

One challenge inherent to current spatial transcriptomic platforms is that each “spot” can capture gene expression information from more than one cell (1-10 cells, characteristically). In a complex tissue such as a healing wound, this often includes cells of different types – particularly within the dermis where fibroblasts, multiple types of immune cells, and nascent blood vessels can be found. As such, to understand our spatial transcriptomics results in the context of our scRNA and scATAC fibroblast data, we needed to account for the contributions of non-fibroblast cells from each Visium spot. This was achieved by first estimating the number of each specific cell type present within individual spots based on the associated histological staining (**Extended Data Fig. 12a-d**). This information was then used to construct single cell gene expression distributions for spots comprised entirely of fibroblasts, keratinocytes, macrophages, neutrophils, or endothelial cells across each timepoint and replicate (**Extended Data Fig. 13**). These sets were then sampled randomly in a Monte Carlo fashion and used to “subtract out” potential contributions from non-fibroblast cells in each fibroblast-containing spot, generating a distribution of 10,000 inferred fibroblast transcriptomes for each Visium spot. These were propagated forward for anchor-based integration to spatially overlaid partial memberships for each of our four scRNA-Clusters (**Fig. 6a**). The resulting partial membership contributions for each spot were then averaged to obtain consensus cluster representations.

We found that the predicted spatial distributions for our scRNA-seq clusters were largely congruent with the transcriptional differences observed earlier between inner and outer cells using our microdissection approach. For example, “mechano-fibrotic” fibroblasts (Cluster 1) became more prominent over time, expanding from the outer to inner wound regions to fill the scar, whereas putative “proliferator” fibroblasts (Cluster 4) were found primarily at the wound center. These “proliferator” fibroblasts were found to be characterized by expression of *Spp1*, a gene closely associated with scar fibrosis (**Fig. 6b, Extended Data Fig. 14a-d**). “Proliferator” fibroblasts also strongly expressed factors that may prime the granulation tissue for keratinocyte migration and proliferation in a paracrine fashion, for example *Mmp9*. Further examining transcriptional programming relative to tissue depth, we observed clear spatial distinctions between the apical and basal regions of the dermis as early as POD 7 and most prominently at POD 14 (**Fig. 6c**). For example, the MMP inhibitor *Timp1* is expressed by fibroblasts in the basal dermis, while *Thbs2*, which mediates cell-matrix interactions, is primarily expressed in the more apical scar region. Furthermore, we observed that chemokines such as *Cxcl5* were preferentially expressed by fibroblasts in the superficial dermis, supporting a role for these cells in regulating inflammation within the wound microenvironment (**Fig. 6c, Extended Data Fig. 14e-f**).

To assess the relative differentiation states of fibroblasts in this system, we applied CytoTRACE to our POD 14 dermal scar data and found that, similar to our scRNA-seq microdissection findings, cells exhibited significantly less transcriptional diversity in the more inner wound regions, further supporting that fibroblasts undergo differentiation as they proliferate inward during tissue repair (**Fig. 6d**). We then applied RNA velocity analysis to these spatial transcriptomic data, comparing the expression of unspliced pre-mRNA and mature spliced mRNA in order to infer directional information for the transcriptional dynamics within the dermis. This approach again predicted a trajectory of differentiation from outer to inner wound regions of the dermal scar (**Fig. 6e, Extended Data Fig. 15a**). Examining genes most highly correlated with RNA velocity, we found *Runx1* to have the strongest alignment along this trajectory of differentiation (**Extended Data Fig. 15b**). *Runx1* has been implicated in myofibroblast differentiation and is known to have cooperative and overlapping roles with its companion gene *Runx2*, which is downstream from and regulated by FAK^32,33,34,35^, again supporting a role for mechanotransduction in driving fibroblast differentiation in wound healing.

## Integrated analysis permits imputation of spatial epigenomic properties

To further explore fibroblast cell fate with spatial resolution, we developed a method to combine our integrated single cell RNA-ATAC framework with Visium in order to impute spatially-informed epigenomes for wound healing fibroblasts (**Fig. 7a**). As described above, we had generated spatial transcriptomic data from unwounded skin and POD 2, 7, and 14 wound tissue following a modified Visium protocol. To extend this analysis to impute spatial epigenomic properties, we used our RNA-ATAC construct to ascribe partial membership values to fibroblasts present within each Visium spot. This was achieved by first attempting to subtract out non-fibroblast contributions, as described above, followed by anchor-based mapping into a higher-dimensional cluster space from our gene integration matrix (**Fig. 7b**). Parameterization was optimized to preserve spatial autocorrelation for the top measured and imputed gene expression distributions within the POD 14 dermis (**Extended Data Fig. 16a-b**). To account for residual contributions from non-fibroblast cells that may remain after our initial subtraction step, we also spiked-in RNA-seq data for keratinocytes, endothelial cells, macrophages, and neutrophils as described above. The resulting putative reference matrix was then used to assign initial partial set memberships for each spatial datapoint using an anchor transfer-based approach. A single-step spatial smoothing filter was applied to this membership space, followed by removal of non-fibroblast contributions and re-normalization. The resulting partial set memberships for each spatial datapoint then allowed us to project higher-order epigenomic features from the scRNA-ATAC data onto these Visium samples (**Extended Data Fig. 17a-d**).

These spatial epigenomic imputations provided a valuable complement to further refine our understanding of the fibroblast biology driving tissue repair. Detailed data analysis is provided in **Fig. 7c-d** and **Extended Data Fig. 18-19** and more broadly summarized below for each time point in the healing process.

Immediately following wound injury, tissue trauma leads to inflammatory cell recruitment, provisional clot formation, and a dermal gap resulting in loss of contact inhibition among local fibroblasts. These fibroblasts are recruited into the wound bed and begin proliferating. Our data suggest that by POD 2, subsets of these cells have differentiated along the wound margin to form a putative “activated-responder” fibroblast subpopulation. Other, less-differentiated and more mechano-sensitive, “mechano-fibrotic” fibroblasts become pre-activated in the deeper dermis at this point, increasing chromatin accessibility for the *Runx1*, which is known to be a primary regulator of mesenchymal progenitor cell proliferation and differentiation^36^.

By POD 7, macrophage-dominated granulation tissue occupies the wound defect, allowing overlying keratinocyte proliferation and re-epithelialization. At this time, “mechano-fibrotic” fibroblasts are engaged in their maximal proliferative activity and begin to differentiate as they finish proliferating toward the wound center and transition to a more “proliferator” subpopulation. These cells are strongly pro-fibrotic and characterized by elevated *Spp1* gene expression and chromatin accessibility (**Fig. 6b**, **Extended Data Fig. 6e**). In parallel, a population of “remodeling” fibroblasts begins to appear in the outer deep dermis (**Fig. 3b**, **Fig. 6a**).

At POD 14, re-epithelialization is complete, and the wound is traditionally considered to be “healed”. However, while keratinocyte activity decreases at this time, there remains a strong immune cell presence maintained by wound fibroblast chemokine secretion to stimulate active fibrosis in the dermal layer (**Fig. 6c**, **Extended Data Fig. 14e-f**). Likewise, while the outer wound “remodeling” and “activated-responder” fibroblast populations appear more quiescent, the “proliferator” fibroblasts enter their most “active” phase at this time, which accelerates fibrosis to a level higher than any prior to complete re-epithelialization, particularly in the inner wound region.

Considering our imputed spatial epigenomics data more globally, we observed that changes to chromatin accessibility frequently preceded downstream changes in gene expression, even within the constraints of our coarse temporal sampling (**Fig. 7c, Extended Data Fig. 18**). For example, we found that the *Runx1* motif, which is downstream from and regulated by FAK mechanotransduction, initially becomes open at POD 2, remains open particularly along the leading wound edge at POD 7, and then begins to decrease in accessibility throughout the nascent scar at POD 14 (**Fig. 7c-d**).

In aggregate, these studies represent a framework for the comprehensive elucidation of wound healing fibroblast phenotypes based on both gene expression and chromatin accessibility – across time, space, and lineage – with unique granularity. Furthermore, these findings allow us to challenge the classical stages of wound healing, typically described as three overlapping phases: inflammation (POD 2), proliferation (POD 7), and remodeling (POD 14)^3^. Based on our findings, we propose a new set of wound healing stages: 1) Early inflammation – in which immune cells are migrating and infiltrating the injury site without proliferation; 2) Re-epithelialization – which includes rapid keratinocyte proliferation across the wound surface, fibroblast recruitment, and macrophage proliferation; and 3) Activated fibrosis – where maximal fibroblast activation is achieved and sustained in a slow asymptotic decay by steady-state inflammatory signaling beneath the “healed” wound (**Extended Data Fig. 20a-b**).

## Conclusions

In this manuscript, we define fibroblast biology throughout the course of wound healing using integrated, single cell multimodal -omics to unravel the spatial, temporal, and functional heterogeneity of these cells. We demonstrate that fibroblasts are activated from tissue-resident cells in response to injury and proliferate polyclonally to fill the wound gap. Furthermore, we demonstrate that fibroblasts undergo spatially-informed differentiation during this process.

Elucidating these relationships required the integration of nascent technologies and data platforms in what is still a rapidly evolving field of multi-omic imputation. To our knowledge, this work represents the first pairing of scRNA-seq analysis with evaluation of single cell chromatin accessibility in the context of tissue repair with spatial resolution. This approach provides a complementary and mutually-informed lens through which we can view these processes, and specifically allowed us to demonstrate that upstream chromatin changes surrounding mechanical signaling elements precede RNA activation and cell proliferation – directly linking tissue force with activation of wound healing fibroblasts.

Furthermore, we were able to identify and characterize putative, functionally-distinct fibroblast subpopulations with divergent transcriptional and epigenomic programs. We designate these four wound healing fibroblast phenotypes as: “mechano-fibrotic”, “activated-responder”, “proliferator”, and “remodeling”. Following wound injury, fibroblast cells are locally-recruited and migrate to the wound. By POD 2, a subset appears to have differentiated to form an “activated-responder” subpopulation, while the remaining outer wound fibroblasts comprise the less differentiated “mechano-fibrotic” cells. These fibroblasts highly express known fibrosis-associated markers such as *Engailed-1* (Rinkevich et al., 2015), *Col1a1* (Xie et al., 2009), *Tgbf2* (Lu et al., 2005), and *Jun* (Wernig et al., 2017). At POD 7, “mechano-fibrotic” cells appear engaged in their maximal proliferative activity and may begin to differentiate in response to mechanotransduction cues as they migrate toward the wound center to become central “proliferator” cells. By POD 14, outer wound “remodeling” and “activated-responder” cells show decreased proliferation and provide support through the secretion of cytokines and coordination of other pro-fibrotic pathways, representing an early steady state maintained by sustained inflammatory signaling within scar tissue.

Taken together, these results illustrate fundamental principles underlying the cellular response to tissue injury. We demonstrate that populations of fibroblasts migrate, proliferate, and differentiate in an adaptive, dynamic response to disruption of their local environment. Understanding the origin, injury-activation, and differentiation trajectories of injury-responsive cells is critical to develop therapeutic strategies to promote tissue repair.

## METHODS

### Animal Models

The following mouse strains were purchased from Jackson Laboratories: Black/6 (C57BL/6J), Actin-Cre^ERT2^ mice (Tg(CAG-cre/Esr1)5Amc/J), eGFP (C57BL/6J-Tg(CAG-EGFP)10sb/J), and FAK^flox^ (B6.129P2(FVB)-*Ptk2^tm^*^1^*^.1Guan^*/J). *α*SMA-Cre^ERT2^ were courtesy of Dr. Ivo Kalajzic, University of Connecticut. Rainbow mice (ROSA26^VT2/GK^^3^) were courtesy of the Weissman Laboratory, Stanford University School of Medicine. All of the mice were genotyped as per manufacturer’s recommendations. Female mice were used for all experiments in this study. Mice were housed at the Stanford University Comparative Medicine Pavilion (CMP) and Research Animal Facility (RAF). The facilities provided light- & temperature-regulated housing. Mice were given rodent chow and water *ad libitum*. A minimum sample size of three animals was used for all experiments (exact numbers for experiments are noted in the figure legends). Animals with appropriate genotypes for a given experiment were randomly allocated to the various experimental conditions. All experiments were completed according to the Stanford University Animal Care and Use Committee standards of care.

### Splinted Excisional Mouse Wound Healing Model

Splinted excisional wounds were created following the protocol outlined by Galiano *et al* ^6^, which was designed to mimic human wound healing kinetics. In brief, mice were anesthetized with isofluorane (Henry Schein Animal Health) at a concentration of 1-2% in oxygen at 3 L/min. The mice were placed in prone position, and dorsal fur was removed. Skin was sterilized with a betadyne wash followed by 70% ethanol. Punch biopsies were used to make 6mm diameter full-thickness dermal wounds. Two dorsal wounds were created on each mouse. A silicone ring was fixed to the dorsal mouse skin using an adhesive and interrupted 6-0 nylon sutures placed around the outer edge of the ring to prevent rapid contraction. A sterile dressing was placed and changed every other day until the wound was harvested. Digital photographs were taken at the time of surgery and every other day at the time of dressing changes.

### Parabiosis

Parabiotic mouse pairs were created as previously described ^37^. In brief, parabiotic pairs consisted of one female wild-type (C57BL/6) mouse and one female eGFP (C57BL/6-Tg(CAG^EGFP^)10sb/J) mouse that were both age-matched and housed together for 2 weeks prior to parabiosis surgery. Mice were anesthetized, shaved, and sterilized as previously described. Matching incisions were made from the base of the elbow joint (olecranon) to the base of the knee joint on corresponding sides of each mouse. The joints and skin were sutured together. Peripheral blood chimerism was determined with flow cytometry two weeks after parabiosis surgery. Wound surgery was performed on the wild-type mouse after systemic circulation was established.

### Liposomal Tamoxifen Induction

Liposomal tamoxifen (LiTMX) was created as per the protocol described by Ransom et al^26^. Briefly, liposomes were applied locally (pulse) to the surface of dorsal mouse wounds to induce Cre recombinase at the time of wounding. Wound surgeries were conducted as described above.

### Tissue Processing

Mouse tissue was fixed in 4% paraformeladehyde (Electron Microscopy Sciences) for 20 hours at 4°C and embedded into paraffin per standard protocols. For cryopreservation, following fixation, specimens were placed in 30% sucrose (Sigma) at 4°C until saturation, followed by OCT at 4°C until saturation and embedded in OCT. Representative tissue specimens were sectioned and stained with hematoxylin and eosin (H&E, Sigma-Aldrich), Picro Sirius Red Stain (Abcam), or Masson’s trichrome (Sigma-Aldrich) per the manufacturer’s protocols.

### Whole Mount

A border of Vaseline was prepared on a Superfrost/Plus microscope slide and the center was filled with mounting medium (BABB clearing reagent or fluoromount-G (SouthernBiotech)). The tissue sample was placed into the reservoir of mounting medium and a cover slip was applied. The whole-mounted samples were stored at 4°C.

### Tissue Clearing

Tissue clearing optimized to preserve expression of endogenous fluorophores as previously described by our laboratory^27^ was pursued on selected Rainbow whole mount and sectioned wound specimens. In brief, for dehydration, tert-butanol (FisherSci) was buffered to a pH 9.5 with triethylamine (FisherSci). Fixed tissue specimens were placed into increasing gradients of tert-butanol (33%, 66%, and 100%) at room temperature for 30 minutes each and then left in 100% tert-butanol overnight. Tert-butanol and benzoic acid:benzyl benzoate (Sigma Aldrich) at a 1:2 ratio were buffered to pH 9.5 with triethylamine (FisherSci). For whole mount samples, tissues were placed in the prepared BABB solution for clearing for 7 hours at room temperature. Cleared whole mount and sectioned tissue specimens were stored in BABB solution at 4°C.

### Confocal Imaging and Analysis

Rainbow mouse tissues were fixed and prepared in the dark to minimize bleaching of endogenous fluorophore expression. Laser scanning confocal microscopy of whole mount and tissue specimens was performed using the Leica WLL TCS SP8 Confocal Laser Scanning Miscroscope (Leica Microsystems) located in the Cell Sciences Imaging Facility (Stanford University, Stanford, CA). Both the 20x and 40x objectives were used for imaging (x20 and x40 HC PL APO IMM CORR CS2, H2O/glycerol/oil, numerical aperture 0.75). Precise excitation and hybrid detection of the Rainbow fluorophores (mCerulean, eGFP, mOrange, and mCherry) was captured. Raw image stacks were imported into either Fiji (NIH) or Imaris (Bitplane/Perkin Elmer) software for further analysis. Z-stacked confocal images were rendered into 3-dimensions. Analysis of clonal volume, area and direction was conducted using the surface and thresholding tools using Imaris software.

### Immunostaining

Cryosections on Superfrost Plus microscope slides (FisherSci) were rehydrated and permeabilized with 0.5% Triton X-100 (Sigma). Tissue sections were then rinsed repeatedly, incubated with 1X Power Block (BioGeneX), and stained with primary antibody for one hour at room temperature. Primary antibodies used for immunostaining included CD45 (D3F8Q) Rabbit mAb (Cell Signaling), Rabbit mAb to alpha Smooth Muscle Actin (Abcam), Rb pAb to Collagen I (Abcam), Rb pAb to Collagen III (Abcam), Rb pAb to Collagen III (Abcam), Specimens were then rinsed repeatedly, stained with a secondary antibody for one hour, rinsed repeatedly, and mounted with ProLong Gold antifade reagent with DAPI. Secondary antibodies used for immunostaining included Goat anti-Rabbit Secondary Antibody Alexa Fluor 488 (Abcam), Goat anti-Rabbit Secondary Antibody Alexa Fluor 647 (Abcam), Donkey Anti-Rat Secondary Antibody Alexa Fluor 647 (Abcam), and Donkey Anti-Rabbit Secondary Antibody Alexa Fluor 555 (Invitrogen). Slides were then coverslipped and imaged using a Lecia DMI6000B inverted microscope or confocal microscopy as described above.

### DAB Immunohistochemistry

Cryosections on Superfrost Plus microscope slides (FisherSci) were rehydrated. H_2_O_2_ was used to quench endogenous peroxidases. Antigen retrieval was conducted using Abcam’s Trypsin Antigen Retrieval kit (Ab970) per the manufacturer’s protocol. Tissue sections were then permeabilized with 0.025% Triton X-100 (Sigma), rinsed, incubated with 1X Power Block (BioGeneX), and stained with primary antibody at 4°C overnight. Primary antibodies used for staining included Phospho-FAK/PTK2 pTyr397 (Invitrogen) and Il6 Monoclonal Antibody (MP5 20F3) (Invitrogen). Tissue sections were then washed, and incubated with secondary antibody at room temperature for one hour. Secondary antibodies used for staining included Horse anti-Rabbit IgG Antibody (H+L) Biotinylated R.T.U. (Vector Laboratories) and Goat anti-Rat IgG Antibody (H+L) Biotinylated R.T.U. (Vector Laboratories). The VECTASTAIN Elite ABC system for avidin/biotin peroxidase (Vector Laboratories) was then applied per the manufacturer’s recommendations. DAB staining was conducted using the BD DAB Substrate Kit (BD 550880) per the manufacturer’s guidelines. Specimens were co-stained with Hematoxylin, rinsed, dehydrated, mounted, and imaged using a brightfield microscope.

### Automated Connective Tissue Analysis

Polarization microscopy images of Picrosirius Red-stained histology specimens were obtained at 40X with a minimum of 10 images per specimen. Images were color deconvoluted to yield separated images of mature (red) and immature (green) connective tissue fibers. These images were then separately denoised, binarized, and quantified across a panel of fiber parameters (length, width, persistence, alignment, etc.), for a total of 294 parameters per image (147 for mature fibers, 147 for immature fibers). Images were quantitatively compared by t-SNE, an algorithm for visualizing high dimensionality data in two dimensions. Individual parameter values were compared between specimen conditions.

### Sample Preparation and Fluorescent-Activated Cell Sorting (FACS) Isolation

Mouse wound tissues were harvested and micro-dissected to separate the outer and inner regions of the wounds. In brief, the wound radius was divided in half, and microdissection was conducted to separate the inner radius and the outer ring based on this division. The wound tissue regions were then placed independently in a dispase-trypsin solution as previously described ^38^ for 30 minutes at 37°C; following this, the epidermis and hypodermis were isolated from the dermal wound scar specimens and discarded. The dermal wound tissues were then minced and digested in DMEM-F12 (GIBCO®) with 0.5mg/ml Liberase^TM^ (Roche) for 30 minutes at 37° in an orbital shaker. The digests were quenched with quench media (DMEM-F12 with 10% Fetal Bovie Serum (FBS, Sigma)), centrifuged at 300 x G for 5 minutes at 4°C, resuspended in quench media, filtered through 100, 70, and 40μm cell strainers (Falcon cell strainer, ThermoFisher), centrifuged once more, and resuspended in FACS buffer.

Cells were counted. Primary antibodies were then applied, and cells were stained in the dark on ice with gentle agitation for 30 minutes. Antibodies against the following cell surface markers primarily or secondarily conjugated to the same fluorophore were used for exclusion of “lineage” cells in order to isolate fibroblasts in an unbiased manner: CD45, CD31, Ter119, Tie2, CD324, and CD326. This approach has been previously validated by our laboratory to isolate mouse wound fibroblasts^29, 39^. Cells were then washed with FACS buffer, centrifuged, and resuspended in FACS buffer. Staining with any secondary antibodies was conducted in the same manner. SYTOX ADvanced Ready Flow Reagent (ThermoFisher) or DAPI (Thermofisher) were used as viability markers. Fibroblasts were isolated using the FACS Aria II system.

For bulk RNA-seq, cells were sorted into chilled lysing reagent under RNA/DNAse-free conditions (Trizol LS, ThermoFisher). For scRNA-seq, cells were bulk sorted by “purity” based on expression of Rainbow colors. These sorted cell aliquots were then individually re-sorted by single cell and index into prepared, 96-well plates containing lysis-buffer. For scATAC-seq, cells were sorted into FACS buffer. mCerulean was arbitrarily selected from among the available Rainbow colors and mCerulean+ wound fibroblasts were used for single-cell sequencing experiments. Flow-cytometry plots shown are representative of at least three independent experiments.

For flow cytometry analysis of cell surface and phosphorylated proteins, a single cell suspension was prepared using manual tissue dispersion rather than enzymatic digestion to preserve phosphorylated signal. Rainbow fibroblasts were isolated as described above and then prepared using the BD Biosciences Cytofix/Cytoperm™ kit according to manufacturer’s instructions. Primary antibodies used for flow cytometric analysis included Phospho-FAK/PTK2 pTyr397 (Invitrogen), DLK1 Monoclonal Antibody (3A10) (Invitrogen), Rat Anti-Mouse Sca1 (Ly-6A/E) (Stem Cell technologies), Rabbit Anti-DPP4 (CD26) (Abcam). Protein expression analysis was conducted using the FACS Aria II system.

FACS gating and data analysis was performed using FlowJo. Gating schemes were established with fluorescence-minus-one controls. Single cells were first gated using FSC and SSC parameters. Dead and lineage-positive (non-fibroblast) cells were then excluded. Gating schemes to quantitate and/or isolate fibroblasts were validated by plating a portion of the sorted cells for morphological visualization.

### Bulk mRNA Sequencing

RNA extraction was performed using Qiagen miRNeasy kit with on column DAnase treatment per the manufacturer’s recommendations. The Clontech Smarter Ultra Low Input RNA kit (Takara Bio) was used to generate cDNA from 150 pg total RNA following the manufacturer’s recommendations. Amplified cDNA was purified using SPRI Ampure Beads (Beckman Coulter) and the quality and quantity was measured using a High Sensitivity DNA chip on the Agilent 2100 Bioanalyzer (Agilent Technologies). cDNA was sheared to an average length of 300 basepairs using a Covaris S2 ultrasonicator (Covaris) and libraries were generated with the Clontech Low Input Library Prep kit (Takara Bio). The samples were uniquely barcoded, pooled, and sequenced on HiSeq (Illumina).

### Bulk mRNA Sequencing Data Analysis

A total of 12 mouse samples were profiled by bulk RNA-sequencing as described above. Raw FASTQ reads were aligned to the GENCODE vM20 reference transcripts (GRCm38.p6) with Salmon^40^ v0.12.0 using the --seqBias, --gcBias, --posBias, --useVBOpt, --rangeFactorizationBins 4, and --validateMappings flags and otherwise default parameters for single-end mapping.

Salmon results were merged into a single gene-level counts matrix using the R package, tximport^41^ v1.4.0. Count normalization and differential gene expression analysis was performed using the DESeq2 v1.22.2 package in R or using Basepair software (www.basepairtech.com). Counts were size-factor normalized using the DESeq function and log2-transformed. Pairwise differential gene expression analysis was performed using the lfcShrink function and indicating type = apeglm, which applies the adaptive t prior shrinkage estimator. As recommended, a threshold of *P*-adjusted < 0.1 was used to define significance for differentially expressed genes.

### Single-cell RNA Sequencing (scRNA-seq)

Mouse wound healing fibroblasts were prepared for PlateSEQ analysis as described above. In brief, mouse dermal scar specimens derived from 4 litter mates were pooled for each timepoint (POD 2, 7, and 14) and divided by wound region (inner and outer). Cells were counted and filtered just prior to loading into the FACS machine. Fibroblasts were FACS-isolated based on expression of individual Rainbow colors, and using an unbiased, lineage-based strategy to isolate fibroblasts independent of cell surface marker expression^29^. The sorted mCerulean+ (arbitrarily selected from among the available Rainbow colors) fibroblast aliquot was then re-, index-sorted into 96-well plates with 4 ul per well of lysis buffer consisting of 1U/ul of Recombinant RNase inhibitor (RRI) (Clontech), 0.1% Triton X-100 (Thermo 85111), 2.5mM dNTP (ThermoFisher), 2.5 uM oligodT30VN (5′AAGCAGTGGTATCAACGCAGAGTACT30VN-3′, IDT). Once sorted, plates were immediately spun down and frozen at -80°C. Single cell RNAseq was performed via the method described by Picelli et al^42^. Lysis buffer plates were thawed on ice, then heated at 72 °C/3 min in a Biorad C1000 Touch thermal cycler to denature RNA. First strand cDNA synthesis was performed in a 10 ul reaction with 100 Units of Clontech’s Smartscribe reverse transcriptase (Cat. No. 639538), 10 Units RRI, 1X First Strand Buffer (Clontech), 5 mM DTT, 1M Betaine (Sigma B0300-5VL), 6mM MgCl_2_, 1 μM Template Switch Oligo (TSO, (5′-AAGCAGTGGTATCAACGCAGAGTACATrGrG+G-3′, Exiqon) at 42°C/90 min, 70°C/5 min. PCR pre-amplification was performed in a 25 ul reaction with 1X Kapa HiFi HotStart, 0.1 uM ISPCR primer (5′-AAGCAGTGGTATCAACGCAGAGT-3′, IDT) at 98°C/3 min, then 25 cycles of 98°C/20 sec, 67 °C/15 sec, 72 °C/6 min, then 72 °C/5 min. Amplified cDNAs were purified by SPRI beads using a Biomek FX automated platform and eluted in 25 ul water, and 2 ul aliquots were run on an Fragment Analyzer High Sensitivity NGS 1-6000 kit for quantitation. Barcoded sequencing libraries were made using the miniaturized Nextera XT protocol of Mora-Castilla et al^43^ in a total volume of 4 ul. Pooled libraries were analyzed on an Agilent Bioanalyzer High Sensitivity DNA chip for qualitative control purposes. cDNA libraries were sequenced on a NextSeq 500 Illumina platform aiming for 50,000 reads per cell.

### scRNA-seq Data Processing

Fastq files for individual cells were converted to BAM format using STAR v2.5.3. Cell barcodes representative of quality cells were delineated from barcodes of apoptotic cells or background RNA based on a threshold of having at least 200 unique transcripts profiled and less than 10% of their transcriptome of mitochondrial origin, resulting in 191 unique single cells, with an average of 550,000 reads per cell. Raw mRNA counts were normalized on a per-cell basis with a scale factor of 10,000 and subsequently natural log transformed with a pseudocount of 1 in R (version 3.6.0) using the Seurat package (version 3.1.1)^44^. Aggregated data was then evaluated using uniform manifold approximation and projection (UMAP) analysis over the first 15 principal components^45^, with n.neighbors = 50, min.dist = 0.75, and repulsion.strength = 1.

### Generation of Characteristic Subpopulation Markers and Enrichment Analysis

Cell-type marker lists were generated with two separate approaches. In the first approach, we employed Seurat’s native *FindMarkers* function with a log fold change threshold of 0.25 using the ROC test to assign predictive power to each gene. However, in order to better account for the mutual information contained within highly correlated predictive genes, we also employed a characteristic direction analysis ^46^. The 50 most highly ranked genes from this analysis for each cluster were used to perform gene set enrichment analysis in a programmatic fashion using EnrichR (version 2.1) ^47^.

### Prediction of Differentiation States

*CytoTRACE* v0.2.1 R package (publicly available at https://cytotrace.stanford.edu)17 was used with default settings to predict differentiation states in scRNA-seq data. Predictions were generated using the CytoTRACE function with a read count matrix as input. Low-dimensional plots for visualizing CytoTRACE and cluster assignments were generated with plotCytoTRACE using UMAP coordinates generated from Seurat. Genes associated with less and more differentiated cells were generated with plotCytoGenes.

### Single-cell ATAC Sequencing (scATAC-seq)

Single cell ATAC-seq was performed following 10x Genomics protocols. In brief, Rainbow mouse wound healing fibroblasts were FACS-isolated using the unbiased, fibroblast-isolation strategy as described above. As for scRNA-seq, mouse dermal scar specimens derived from 4 litter mates were pooled for each timepoint (POD 2, 7, and 14) and divided by wound region (inner and outer). As for scRNA-seq, the sorted mCerulean+ (arbitrarily selected from among the available Rainbow colors) fibroblast aliquot was selected for further analysis. For nuclei isolation, 10x genomics protocol CG000169 Rev D was followed using the low cell input modifications. Transposition, GEM generation and barcoding, post GEM Incubation Cleanup, library construction, and quantification was conducted following the 10x genomics protocol CG000169 Rev D. Pooled libraries were analyzed on an Agilent Bioanalyzer High Sensitivity DNA chip for qualitative control purposes. cDNA libraries were sequenced on a NextSeq 500 Illumina platform. In total, we generated scATAC-seq profiles from 5,353 cells, which yielded on average 13.0 × 10^3 unique fragments mapping to the nuclear genome.

### scATAC-seq Data Processing and Analysis

Raw base call (BCL) files were demultiplexed to fastq files using the 10x Genomics Cell Ranger tool *cellranger-atac mkfastq*. These files were then aligned to the mouse genome (mm10) using *cellranger-atac count* with default parameters. Downstream analysis of scATAC-seq data were performed using ArchR, a novel tool developed by our collaborators (Granja et al., 2020). Arrow files were created for each sample using TSS and frag filters of 4 and 1000, respectively. Doublets were filtered using k = 10 pseudodoublets embedded in UMAP space, as previously described (Granja et al., 2020). Dimensionality reduction was achived using ArchR’s implementation of latent semantic indexing (LSI). Initial clustering was achieved using the native *addClusters* function with resolution = 0.1, method = Seurat, reducedDims = IterativeLSI. Single cell embeddings were then generated using *addUMAP* with nNeighbors = 40, minDist = 0.5, metric = “cosine”, and marker genes were identified using *getMarkerFeatures* with bias = c(”TSSEnrichment”, “log10(nFrags)”), testMethod = “wilcoxon”.

### Integration of scRNA-seq and scATAC-seq Data

Single cell ATAC-seq data were integrated with scRNA-seq data using ArchR. Cells from scATAC-seq are directly aligned with cells from scRNA-seq by comparing the scATAC-seq gene score matrix with the scRNA-seq gene expression matrix. This alignment is performed using the *FindTransferAnchors()* function from Seurat, which allows for the alignment of data across two datasets. This cross-platform linkage is performed serially in an unconstrained and constrained fashion. Pseudo-scRna-seq profiles are then generated for each single ATAC cell using the native ArchR *addGeneIntegrationMatrix()* function with default parameters, and scATAC-seq clusters were then labeled with scRNA-seq information. Pseudo-bulk replicates were then generated using ArchR’s *addGroupCoverages()*, permitting peak calling with ArchR’s native TileMatrix algorithm. Motif enrichment calculations were performed using *addMotifAnnotations* and p*eakAnnoEnrichment* with cutOff = “FDR <= 0.1 & Log2FC >= 0.5”.

### GREAT Analysis

Genomic Regions Enrichment of Annotations Tool (GREAT) analysis was performed programmatically in R (rGREAT v1.18.0) to assess for enrichment of cis-regulatory regions ^30^. This was performed separately for the initial six ATAC-defined clusters and the subsequent four clusters defined based on integrated scRNA-seq analysis. In each case, this was performed iteratively for each cluster using pairwise comparisons between peak sets for that cluster against a background containing the union of all peak sets from the dataset (including those for that cluster).

### Deconvolution of Bulk RNA-seq Profiles

CIBERSORTx^19^ was used to deconvolve cell type abundances from bulk RNA-seq profiles of fibroblasts isolated from untreated wounds at POD7 (n = 4) and POD14 (n = 3) and FAK-inhibitor-treated wounds at POD14 (n = 4). Default parameters from the web toolkit (http://cibersortx.stanford.edu/) were used to generate a custom signature matrix using a single cell reference matrix file consisting of raw mRNA counts for each cell in the scRNA-seq dataset. Each scRNA cell was re-labeled using Seurat’s anchor-transfer mapping algorithm to the integrated ArchR scRNA-ATAC reference profiles: “ArchR-Cluster 1”, “ArchR-Cluster 2”, “ArchR-Cluster 3”, and “ArchR-Cluster 4” subpopulations as defined in **Fig. 3b**. Imputation of cell fractions was then performed using this matrix in conjunction with a mixture file of bulk RNA-seq samples, using S-mode batch correction without quantile normalization.

### Spatial Transcriptomics

Wound specimens were rapidly harvested and flash frozen in OCT. Using the Visium Tissue Optimization Slide & Reagent kit, permeabilization time was optimized at a thickness of 10um per section and 37 minutes for mouse cutaneous scar tissue. Wound tissues were cryosectioned at - 20 degrees onto gene expression slides. The Gene Expression Slide & Reagent kit was followed per protocol, and used to produce sequencing libraries. The libraries were then sequenced using NextSeq (Illumina), and Bcl files were demultiplexed.

Raw FASTQ files and histology images were processed by sample with the Space Ranger software, which uses STAR v.2.5.1b (Dobin et al., 2013) for genome alignment, against the Cell Ranger mm10 reference genome, available at: http://cf.10xgenomics.com/supp/cell-exp/. Raw spaceranger output files for each sample were read into a Seurat class object in R using Seurat’s Load10X in a manner that kept them paired with the low resolution histology images for visualization purposes. This included information such as the number of estimated cell counts, the sum of UMIs per spot, number of expressed genes per spot, and graph-based clustering results. We did not drop any spots given the spatial pattern they presented. Data were normalized using the SCTransform with default parameters. Principal component analysis was performed on the normalized dataset, and the top 30 components were used for neighbor finding, Louvain-based cluster analysis, and UMAP dimension reduction and embedding.

For the delineation of cell type contributions to each Visium “spot”, high resolution histology images (H&E) were loaded and each spot was counted for the number of a given cell type in that spot. This was performed by co-authors who were not involved in the Visium data analysis. A maximum of 9 cells of a given type were used as the limit for a given spot. Spots consisting entirely of one cell type, based on these annotations, were used to construct sets of “pure” keratinocyte, endothelial cell, macrophage, and “neutrophil” (including all non-macrophage immune cells) transcriptional profiles. We then sampled from these in silico population in a Monte Carlo fashion using 10,000 iterations to “subtract out” the contribution of non-fibroblast cells from each fibroblast-containing spot in a linear fashion. Negative gene expression values were zeroed out at a per-spot level following all subtractions, representing an asymmetrical bias for spots based on the number of non-fibroblast cells, which we found to counterbalance the assumption of linear added effects. These values were propagated forward for anchor-based integration to spatially overlaid partial memberships for each of our four clusters. The resulting partial membership contributions for each spot were then averaged to obtain consensus cluster representations

RNA velocity analysis was performed using scVelo,^48^ using a likelihood-based dynamical model to solve the full transcriptional dynamics of mRNA splicing kinetics. Whereas the originally described steady-state model assumes that all genes share an equal splicing rate ^49^, the likelihood-based dynamical model of scVelo generalizes RNA velocity analysis to transient cell states and non-stationary subpopulations.

### Imputed Spatial Epigenomics

In order to impute spatial epigenomic properties from our Visium datasets, we first re-partitioned our scRNA-ATAC construct with higher cluster resolution in order to better capture differences in the time and space phenotypes represented by the underling scRNA and scATAC data. This parameterization was optimized using our POD 14 spatial transcriptomic data, which had the clearest delineation of dermal margins and highest representation of fibroblasts. We first defined an “outer” ↑→ “inner” vector along the scar dermis and then computed the spatial auto-correlation for each gene along this vector using six neighbors for each Visium spot (**Extended Data Fig. 16a**) ^13^. We considered the top 100 most highly auto-correlated genes to have spatial significance in our original dataset (**Extended Data Fig. 16b**). We then iteratively re-partitioned our multimodal scRNA-ATAC fibroblast data using ArchR’s native Louvain clustering algorithm while varying the resolution parameter from 0.5 to 4.0 at increments of 0.1. Each of the resulting cluster configurations was used to generate partial membership profiles for each Visium spot using the anchor-based transfer method described above, after which the Gene Integration Matrix values associated with each ArchR cluster were projected onto each Visium spot in accordance with the each cluster’s partial spot membership. Preservation of auto-correlation for the 100 genes determined above was used as an optimization metric, and we selected a resolution parameter of 2.2, which produced twenty clusters, each representing between 27 and 552 cell equivalents. Gene integration matrix distributions, informed by both modalities, were then extracted for each partition and subjected to SCT transformation. “Spike-in” scRNA-seq data for keratinocytes, endothelial cells, macrophages, and neutrophils were generated as described above and augmented to the initial 20 RNA-ATAC-defined clusters following a similar SCT transformation. The resulting putative single cell gene expression reference matrix (comprising 24 cell-types) was then used to assign initial partial set memberships for each spatial transcriptomic datapoint using an anchor transfer-based approach. Given that the Visium technology captures multiple cells within each 55 um spot, and discretizes all of their location to the spatial center of that spot, we applied a single-step spatial smoothing function to incorporate fractional membership contributions from each of the six neighboring spots. This is achieved by updating each partial membership value *mi* for each *spotj* in the Visium slide:

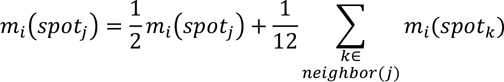

Partial memberships to non-fibroblast clusters were subsequently removed from each spot, and the remaining partial membership values for each fibroblast cluster were normalized to enforce a membership sum of 1 for each spot:

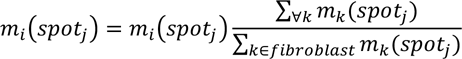

The resulting partial set memberships for each spatial datapoint were then treated as a topological vector space, onto which epigenomic peak, motif, and binding activity from the twenty scRNA-ATAC partitions can be projected.

To examine the spatial dependence of individual genes along the out-to-inner wound axis, spatial lag vectors were constructed for POD 14 Visium samples using a 3-spot diameter trajectory as shown in (**Extended Data Fig. 16a**)^13^, and Pearson correlations calculated for imputed gene scores along either radial direction. The top 200 genes from each group were used to perform enrichment analysis against the Gene Ontology database. Functional networks of the most highly enriched gene sets were then generated using the clusterProfiler package in R ^50^.

### Statistical Analysis

Non-omics statistical analyses were performed using the software GraphPad Prism v.6 (unless otherwise noted). Results are expressed as absolute numbers, percentages, fractions, or mean +/-standard deviation (unless otherwise noted). Unpaired *t*-test assuming two-tailed distribution or one-way analysis of variance (ANOVA) and post hoc Tukey correction were used to compare groups where relevant. *P* < 0.05 was considered statistically significant. All -omics statistical analyses were performed in R (version 3.6.0) as described above.

## Acknowledgements

The authors would like to acknowledge Sopheak Sim at the Stanford Functional Genomics Facility (SFGF), Krista Hennig and the Stanford Genome Sequencing Service Center (GSSC), Daphne Cooper and Christina Chiu at 10x Genomics for assistance with gene expression experiments, Jonathan Mulholland and Kitty Lee and the Beckman Cell Sciences Imaging Facility (CSIF), Catherine Carswell-Crumpton and the Stanford Stem Cell FACS Core, Michael Lopez for his assistance with flow cytometry experiments, Chloe Steen for her assistance CIBERSORTx, Alessandra Moore, Eliza Foley, and Emma Briger for their assistance with animal experiments and processing of specimens, and George Yang, Charles C Chan, Sidd Menon, and Natalina Quarto for their invaluable advice regarding experimental design and manuscript preparation.

Sequencing data for this project was generated on an Illumina NextSeq 500 and an Illumina HiSeq 4000 that were purchased with funds from the NIH under award number S10OD018220. The confocal imaging data for this project was obtained using the Leica SP8 that was purchased with funds from the NIH under award number 1S10OD01058001A1.

Funding sources for this project include American College of Surgeons Resident Research Scholarships (D.S.F.), NIH NCI Postdoctoral Individual National Research Service Award 1F32CA239312-01A1 (D.S.F), the Advanced Residency Training at Stanford (ARTS) program (D.S.F.), the Stanford Transplant and Tissue Engineering Center of Excellence Fund (R.E.J.), the Gunn/Olivier Fund, the California Institute for Regenerative Medicine, the Hagey Laboratory for Pediatric Regenerative Medicine, Stinehart/Reed Foundation, Goldman Sachs Foundation administered through the Stanford Cancer Institute (J.A.N., D.S.F., M.T.L.), NIH 1R01GM116892 (M.T.L), NIH 1R01GM136659 (M.T.L), 5U01DK119094 (M.J., G.C.G.), NIH R00CA187192-03 (A.M.N.), K08DE024269 (D.C.W.), the Child Health Research Institute (CHRI) at Stanford University (D.C.W.), NIH RM1-HG007735 (H.Y.C.), and the Scleroderma Research Foundation (H.Y.C.). H.Y.C. is an Investigator of the Howard Hughes Medical Institute.

## Author Contributions

D.S.F. conceived of the study, performed experiments, analyzed data, produced figures, and wrote and edited the manuscript; M.J. conceived of the study, contributed to experimental design, analyzed data, developed and implemented the multimodal data integration methods, produced figures, and wrote and edited the manuscript; M.S.C. contributed to experimental design, performed experiments, analyzed data, contributed to figures, and wrote the manuscript; K.Y., G.G., D.H., and K.T. contributed to experimental design, performed experiments, analyzed sequencing data, and contributed to figures; A.S., A.T.N., A.R.B., R.C.R., S.M., A.L.T., R.E.J., O.dS., W.T.L., C.D.M., H.dJP., K.C., and M.H. performed experiments, analyzed data, and contributed to figures; D.H. and J.C. conducting sequencing experiments and guided analysis; D.C.W., J.A.N, G.W., G.C.G., A.M.N., H.Y.C., and M.T.L. conceived of the study, guided experimental design, guided data analysis, and wrote and edited the manuscript. All authors discussed the work, reviewed, and approved the final manuscript.

## Declaration of Interests

H.Y.C. is a co-founder of Accent Therapeutics, Boundless Bio, and an advisor of 10x Genomics, Arsenal Biosciences, and Spring Discovery.

## Notes

### Competing Interest Statement

H.Y.C. is a co-founder of Accent Therapeutics, Boundless Bio, and an advisor to 10x Genomics, Arsenal Biosciences, and Spring Discovery.

