## Supplemental Information for "Integrated spatial multi-omics reveals fibroblast fate during tissue repair"

### EXTENDED DATA FIGURE LEGENDS

#### Extended Data Figure 1.

**a,** Representative stitched image of a complete mouse wound specimen on cross section immuno-stained for expression of PDGFRA, COL1, and COL3, with DAPI for nuclei. White dotted lines indicate wound scar edge. Solid box indicates zoom of region indicated on figure panel. *n* = 3 per condition.

**b,** Schematic illustrating how tissue-resident fibroblasts (**left panel**) are activated in response to wound injury (**middle panel**) and contribute to scar fibrosis (**right panel**).

**c,** Representative photograph showing a fresh wound prepared using the stented dorsal wound healing model employed in this study, which limits contraction of the panniculus carnosus and thereby mimics human wound healing kinetics [6]. Black dotted line circumscribes the wound edge.

**d,** Schematic of the mouse parabiosis model used, with dorsal wounds created on the wild-type (Black/6) mouse parabiosed to an eGFP mouse after shared blood supply has been established between the parabionts.

**e,** Representative FACS-plots showing analysis of blood taken from a wild-type (Black6) parabiont 2 weeks following parabiosis to an eGFP mice. Establishment of a shared blood supply is indicated by the presence of circulating GFP+ cells (green box).

**f,** Representative histologic sections from wounds made on the wild-type parabiont and harvested at post-operative day (POD) 7 (**left panels**) and POD 14 (**right panels**), demonstrating systemically-derived (GFP+) cells within the scar area; the vast majority of these cells co-stain for CD45. DAPI (blue) identifies cell nuclei. Thick white dotted lines indicate dermal scar area, and thin white dotted lines highlight individual GFP+ DAPI+ cells. Right panels correspond to the sub-regions outlined by white solid boxes. *n* = 3 per condition per timepoint. Scale bar = 25μm.

**Extended Data Figure 2.**

**a,** PCA plot of bulk RNA-seq of wound specimens harvested at POD 7 showing clustering of inner wound region fibroblast gene expression as distinct from the corresponding outer wound region fibroblasts and uninjured control skin fibroblasts.

**b,** Volcano plot highlighting genes of interest (pro-fibrotic and mechanotransduction-related) upregulated in wound conditions compared with fibroblasts isolated from uninjured control skin from bulk RNA-seq displayed above.

**c,** Selected heatmap displaying hierarchical clustering for inner wound region fibroblasts compared with outer and uninjured control skin and highlighting relative expression of specific fibrosis- and mechanotransduction-related genes among the aforementioned fibroblast samples. Scale at right represents log fold-change.

**Extended Data Figure 3.**

**a,** Representative FACS-plots showing isolation of Rainbow wound fibroblasts. Images at right show plated, freshly-isolated Rainbow fibroblasts for morphological and fluorophore-expression validation. Rainbow colors as labelled in figure.

**b,** Flow cytometry analysis of FACS-isolated Rainbow wound fibroblasts for expression of cell surface markers associated with skin fibroblast subpopulations. $n = 6$.

**c,** Flow cytometry analysis for expression of cell surface markers associated with skin fibroblast subpopulations of FACS-isolated Rainbow wound fibroblasts separated into inner and outer wound regions. $n = 3$.

**Extended Data Figure 4.**

3D embedding of CytoTRACE data shown in **Fig. 2f** further delineates cells based on predicted differentiation capacity.

**Extended Data Figure 5.**

**a,** Overlay of transcription start site (TSS) enrichment and fragment enrichment for scATAC-seq samples (**top panels**). Violin plots showing TSS and fragment enrichment for each sample type analyzed by scATAC-seq (**bottom panels**). POD = post-operative day, "O" = Outer wound region, "I" = Inner wound region.

**b,** Breakdowns of isolation time point and inner versus outer region status are provided for each scATAC-seq cluster. Y-axis displays cell fraction of cluster total for each of the 6 scATAC-seq conditions; individual fraction values are noted above each bar.

**c,** Heatmap with hierarchical clustering highlighting relationships between scATAC-seq specimens based on phenotype and cluster assignment. POD = post-operative day, I = inner, O = outer.

**d,** Mapping of scATAC-only defined clusters to integrated scRNA-ATAC clusters.

**Extended Data Figure 6.**

**a,** UMAP embedding of integrated scRNA-seq and scATAC-seq showing four multimodal fibroblast clusters, provided for reference.

**b,** Heatmap of cluster peaks shown in (**a**) demonstrating aggregate heterogeneity in chromatin accessibility.

**c,** Motif heatmap highlighting key gene loci differentially open or closed in putative fibroblast subpopulations.

**d,** Plots showing integrated ArchR peaks per cluster at top of each plot, and association between the peaks with fibrosis and mechanotransduction-related genes (Peak2GeneLinks) at the bottom of each plot. FAK (*Ptk2*) is shown in top left panel. Pale orange shading highlights differentially expressed peaks across the ArchR-Clusters. All highlighted peaks demonstrated statistically significant differential expression in at least one pairwise comparison with FDR < 0.1 and FC >= 2.

**e,** Feature plots showing integrated ArchR chromatin accessibility and gene expression data for fibrosis- and mechanotransduction-related gene elements of interest.

**Extended Data Figure 7.**

**a,** Additional tracking plots showing integrated ArchR peaks per cluster at top of each plot, and association between the peaks with fibrosis and mechanotransduction-related genes (Peak2GeneLinks) at the bottom of each plot. Pale orange shading highlights differentially expressed peaks across the ArchR-Clusters. All highlighted peaks demonstrated statistically significant differential expression in at least one pairwise comparison with FDR $< 0.1$ and FC $>= 2$.

**b,** Additional feature plots showing integrated ArchR chromatin accessibility and gene expression data for the ArchR-Clusters from A in terms of fibrosis- and mechanotransduction-related genes of interest.

**Extended Data Figure 8.**

Notable transcription factor (TF) footprinting based on accessibility at a given motif across all identified ATAC peaks, from the integrated ArchR dataset based on pseudo-bulk peak calls. Top row highlights TF footprinting plots relevant for ArchR-Cluster 1, second row for ArchR-Cluster 2, third row for ArchR-Cluster 3, and fourth row for ArchR-Cluster 4.

**Extended Data Figure 9.**

**a,** GREAT analysis of ArchR Cluster data showing enrichment for specific gene sets of interest.

**b,** ArchR-generated pseudotime plot using standard UMAP embedding (left panel), and heatmaps showing Gene Score values imputed from chromatin peak accessibility (middle panels) and Gene Integration values imputed from integration of scRNA-seq data (right panels) relative to pseudotime.

**Extended Data Figure 10.**

**a,** Representative DAB staining for expression of pFAK in vehicle control-treated wounds (**left top panel**) compared with $FAK^{fl/+}$ (**left middle panel**) or FAKi-treated specimens (**left bottom panel**). $n$ = 3 per condition. Quantitation showing fraction of pFAK+ cells per HPF per condition (**right panel**)

**b,** Representative photographs of healing wounds taken every other day over 14 days for the following conditions: vehicle control (**top sub-panel, blue**), FAKi (**second sub-panel, purple**), $FAK^{fl/+}$ (**third sub-panel, red**), and $FAK^{fl/fl}$ (**bottom sub-panel, grey**). Dotted black lines indicate open wound area. Solid yellow line highlights the scar area within healed wounds at POD 14.

**c,** Quantitation of wound healing data from left panels shows no significant difference in rate of wound re-epithelialization among groups.

**d,** H&E (**top panels**) and trichrome (**bottom panels**) of stitched images of complete wound specimens for vehicle control (top panels) compared with FAKi-treated wounds at POD 14. White dotted lines indicated edge of wound scar area. $n$ = 5.

**e,** Representative picrosirius red staining of wound specimens: uninjured skin (**top left**), vehicle-control wound (**top right**), FAKi-treated wound (**bottom left**), and $FAK^{fl/+}$ wound (**bottom right**). $n$ = 3 per condition. Scale bars = 50$\mu$m.

**f,** Automated connective tissue analysis of polarization microscopy images of Picrosirius Red-stained histology specimens shown in (**e**). The connective tissue ultrastructure in FAKi-treated or from $FAK^{fl/+}$ specimens is more similar to unwounded skin than vehicle-control (DMSO) wounds on t-SNE analysis. Sample types highlighted with colored dotted lines (**left panel**). Quantitative analysis of mean red (**middle panel**) and green (**right panel**) fiber intensity for Control (injured) skin, and DMSO (vehicle control), $FAK^{fl/+}$, and FAKi-treated wounds. Statistically significant differences are seen with FAKi-treatment in terms of both fiber types.

**Extended Data Figure 11.**

**a,** PCA-analysis of bulk RNA-seq data for mouse fibroblasts isolated from wounds treated with FAKi versus control and separated in terms of inner and outer wound regions at POD 14.

**b,** Sample-level heatmap with hierarchical clustering of bulk RNA-seq data from inner and outer wound regions with or without FAKi treatment.

**c,** EnrichR analysis for bulk RNA-seq data comparing FAKi-treated wound fibroblasts with vehicle-control wounds. Left panels show gene pathways down with FAKi, right panel shows gene pathways up with FAKi. Top three panels show the 20 most significant GO terms for each cell condition (Biological function, Cellular function, Molecular function). Bottom panels show the top 20 pathways from the BioPlanet 2019 database. All enrichment analyses were conducted using the top 100 most significant genes based on adjusted p-value.

Data represent means +/- S.D. **P* = <0.0001, ***P* = <0.0001, ****P* = 0.0003, *****P* = <0.0001

**Extended Data Figure 12.**

**a-d,** Manual annotation of cell types within Visium spots based on H&E histology for (**a**) POD 0 (unwounded), (**b**) POD 2, (**c**) POD 7, and (**d**) POD14. Panels to the right of (**c**) show zoomed views for an arbitrary wound region, with the second zoomed panel representing a single Visium spot. Neutrophils are circled with green (with characteristic multi-lobed nuclei) in the lumen on a small vein with endothelial cells circled with orange.

**Extended Data Figure 13.**

Projection of manual cell type annotations onto Visium slides across all four time points. In this context we reference “other immune” cells, rather than neutrophils (as initially ascribed in **Extended Data Fig. 12**), to reinforce that these assessments are simply estimations and as such are subject to considerable variability.

**Extended Data Figure 14.**

**a-f,** Visium plots showing POD 0, 2, 7, and 14 (top to bottom) wound sections. Expression of *Ctss*, which is a known mechanoresponsive gene[51], closely aligns with *Spp1*+ *Mmp9*+ Cluster 4 fibroblasts (see **Fig. 6**) along the closing edge at POD7 (**a**). *Postn* expression appears along the dermis later over the wound healing course at POD7 and then localizes to the suprabasal region at POD 14 (**b**), whereas *Tnc* expression appears more in the inner wound at POD7 and then is expressed diffusely across wound fibroblasts at POD14 (**c**). *Col1a1* expression is very prominent across the scar over the course of wound healing (**d**). Expression of *Ccl9* and *Cxcl1* is prominent in the wound dermis particularly at POD14 highlighting how fibroblasts regulate immune cells within the wound microenvironment, similar to *Cxcl5* (see **Fig. 6**) (**e-f**).

**Extended Data Figure 15.**

**a,** RNA-velocity analysis applied to POD 14 Visium data, with spots colored according to tissue layer.

**b,** RNA-velocity data for *Runx1*, the gene whose marginal expression was maximally correlated with the predicted differentiation arc. Plot displaying ratio of unspliced to spliced mRNA at top, velocity data at middle, and expression data at bottom.

**Extended Data Figure 16.**

**a,** “Outer” ←→ “inner” vector along central dermis.

**b,** Spatial lag vector, encompassing six hexagonal "neighbors” for each Visium spot.

**Extended Data Figure 17.**

**a-d,** Overlay of scRNA-ATAC partial probability memberships for each of the twenty partitions onto Visium histology sections at (**a**) POD 0 (unwounded skin), (**b**) POD 2, (**c**) POD 7, and (**d**) POD 14.

**Extended Data Figure 18.**

**a-c,** Visium plots showing POD 0, 2, 7, and 14 (top to bottom) wound sections, imputed spatial epigenomics. **(a)** For housekeeping genes such as *Col12a1* (top panel), gene imputed matrix (GIM) correlates with gene score matrix (GSM) epigenomic data and is fairly stable over space and time. **(b)** However, for the profibrotic gene *Fos*, which is very active within wound fibroblasts, GSM data shows opening at the *Fos* motif along the outer wound at POD2, which yields strong gene expression across wound fibroblasts at POD 7. **(c)** Visium motif deviation plots showing POD 0, 2, 7, and 14 (top to bottom) for *Fos* shows significant openness proximal to the *Fos* motif across the wound area at POD7, likely preceeding gene expression in this area which is present at POD 14.

**Extended Data Figure 19.**

**a-b,** Using the spatial lag vector described in **Extended Data Fig. 16**, Pearson correlations were calculated for epigenomically-imputed chromatin accessibility on a per-gene basis along either radial direction. The most highly correlated genes from each direction (**b**) were then used to generate functional enrichment plots using the Gene Ontology (GO) database (**a-b**).

**Extended Data Figure 20.**

**a-b,** The findings presented here challenge the classical stages of wound healing (**a**), typically described as three largely discrete phases: inflammation (POD 2), proliferation (POD 7), and remodeling (POD 14). We propose a new framework for viewing the stages of wound healing: 1) Early inflammation, 2) Re-epithelialization, and 3) Activated fibrosis – where maximal fibroblast activation is achieved and sustained by steady-state inflammatory signaling beneath the "healed" wound (**b**).

### Extended Data Figure 1

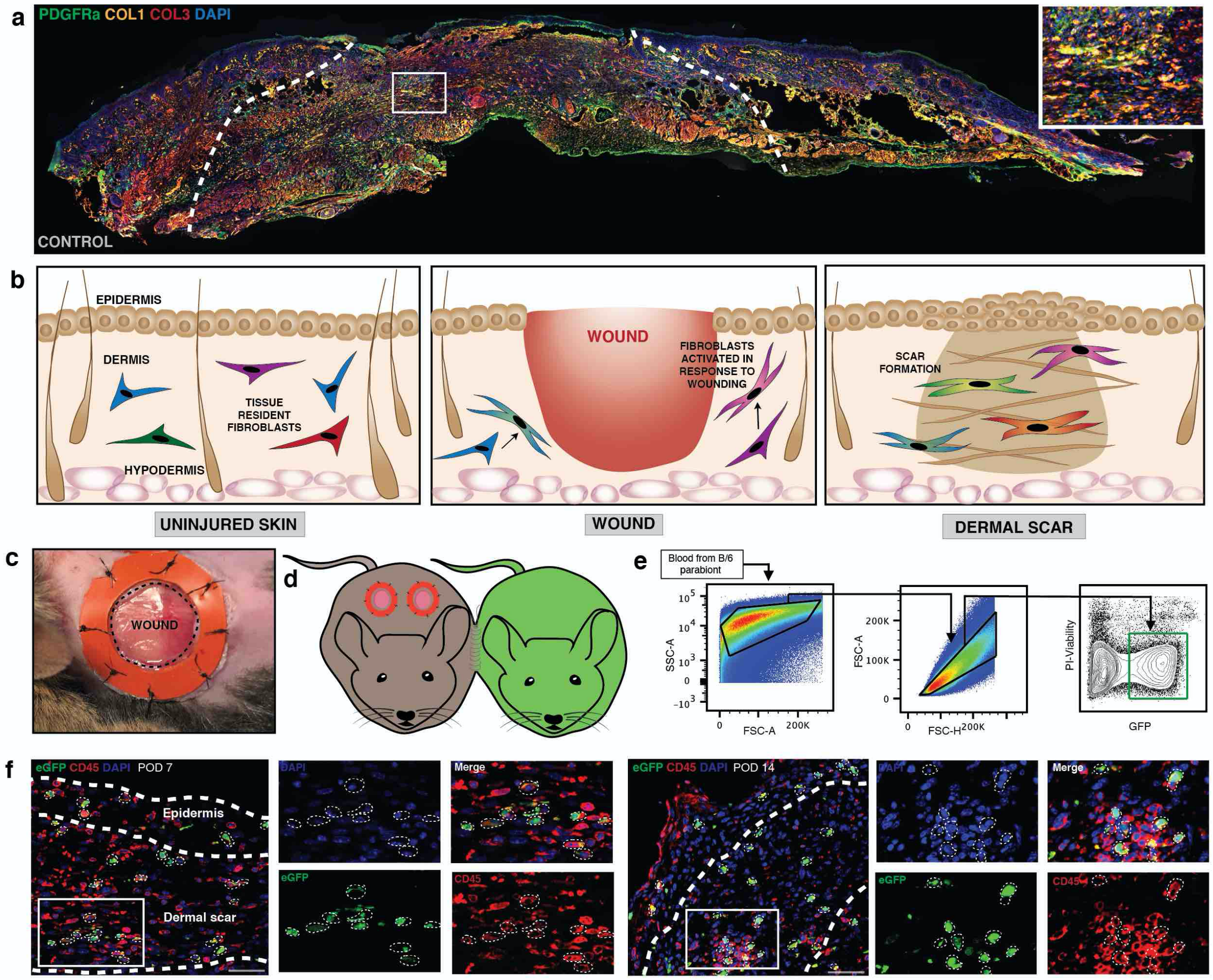

Extended Data Figure 2

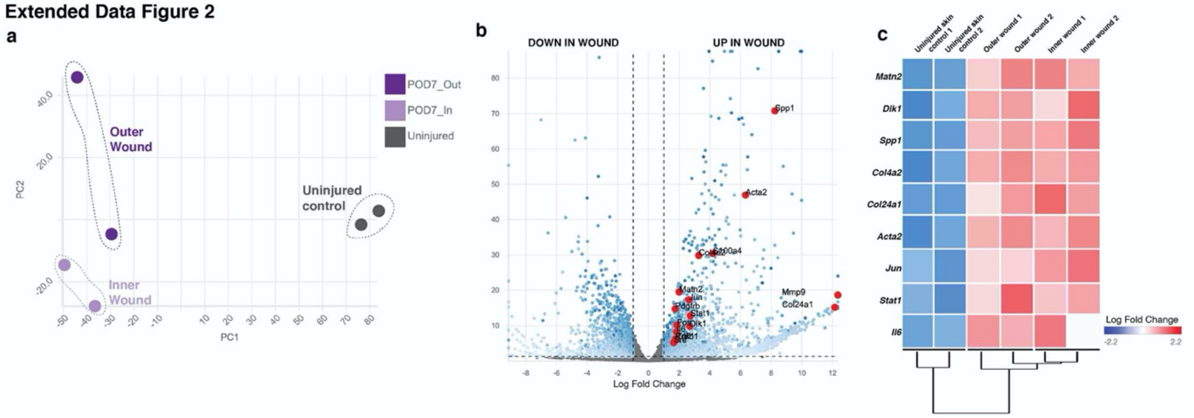

**Extended Data Figure 3**

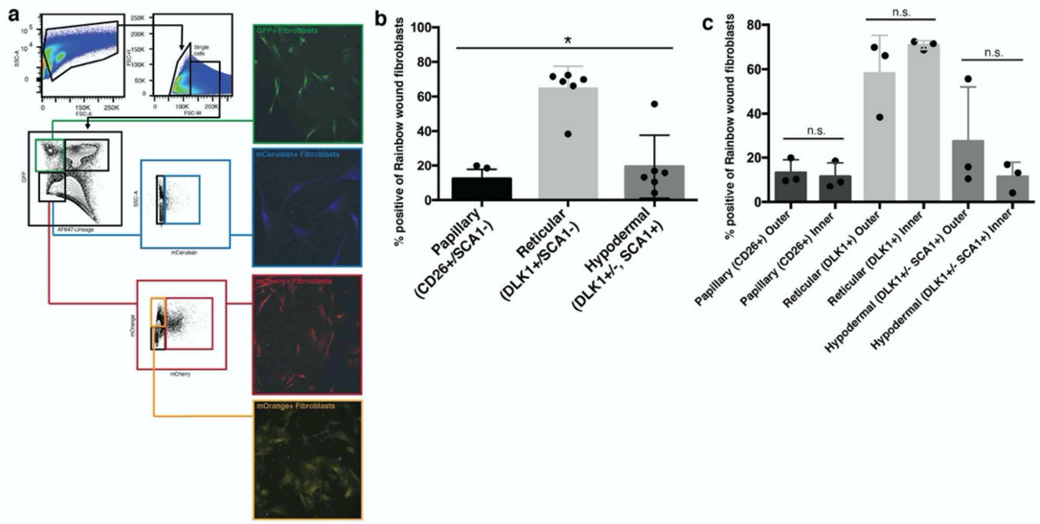

### Extended Data Figure 4

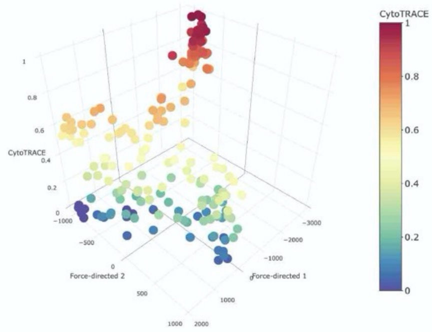

### Extended Data Figure 5

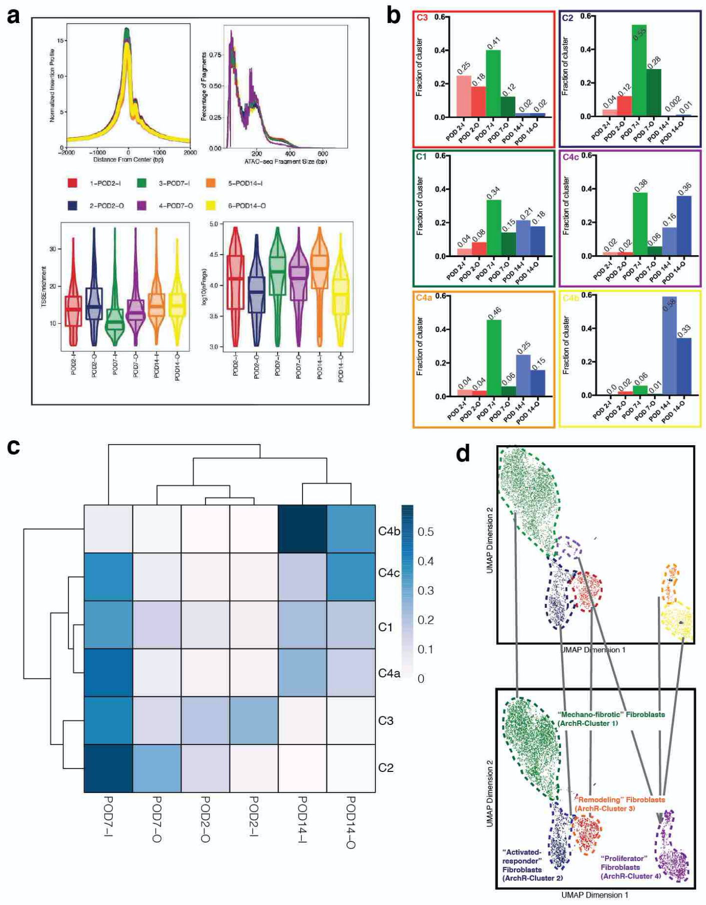

### Extended Data Fig. 6

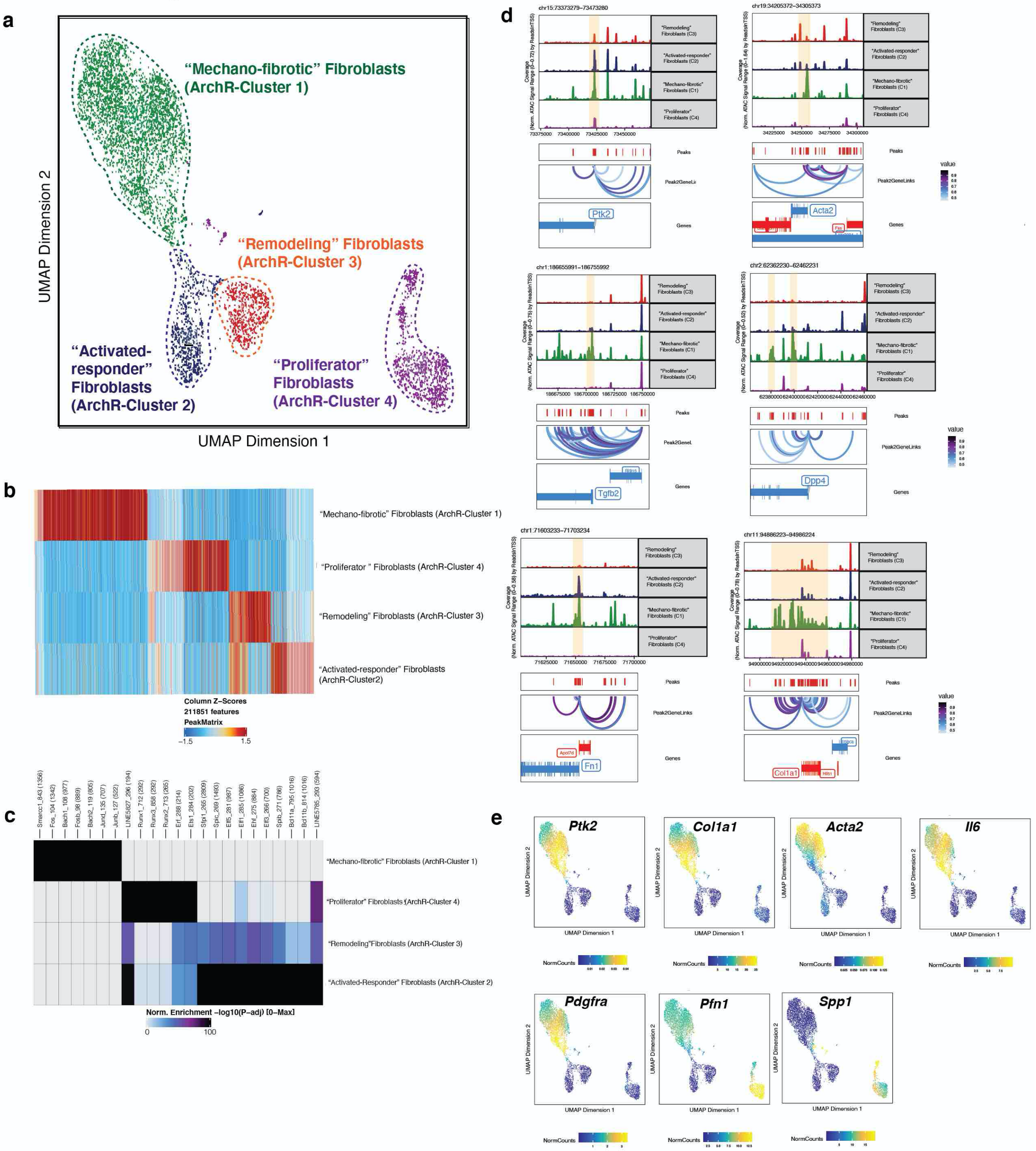

### Extended Data Figure 7

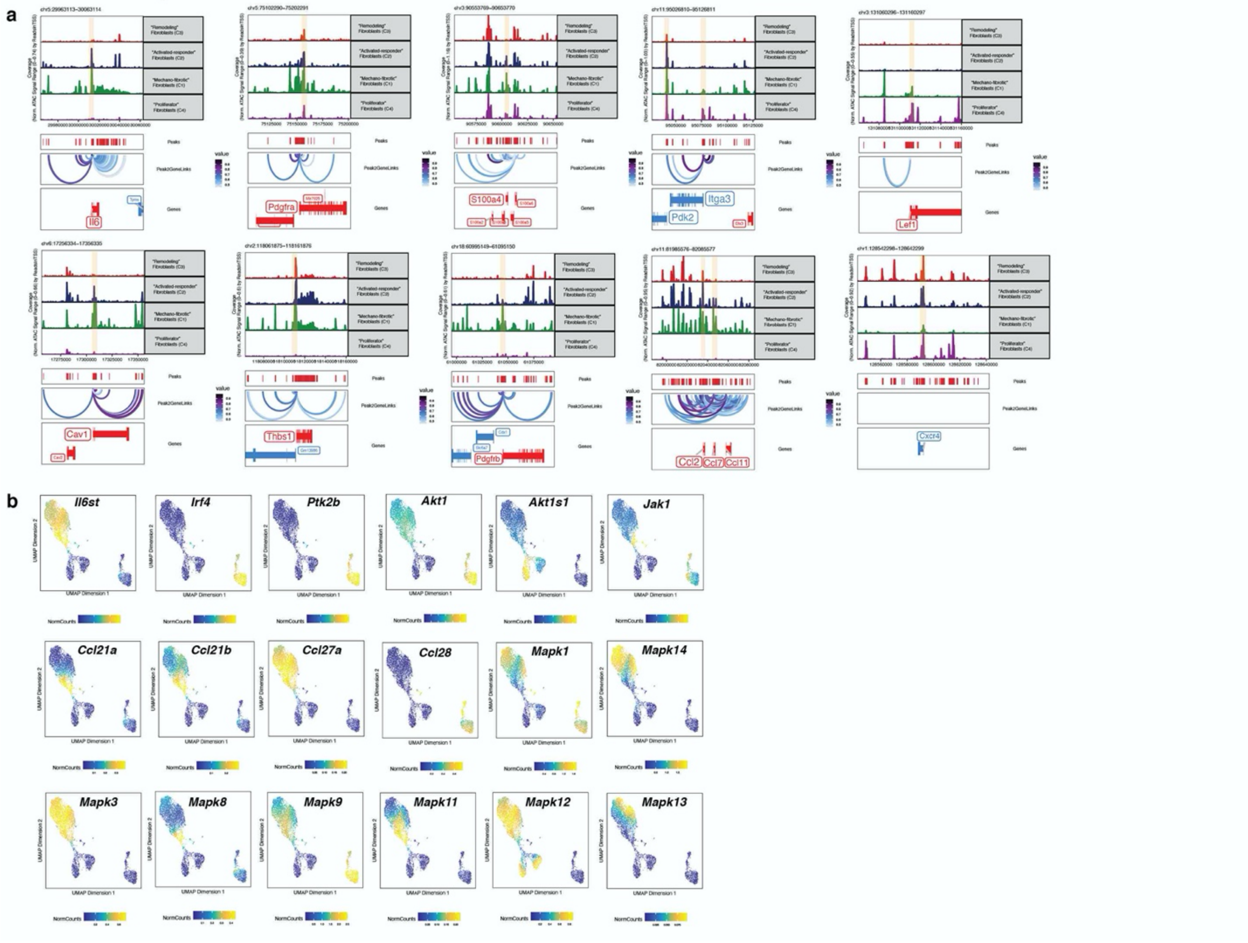

### Extended Data Figure 8

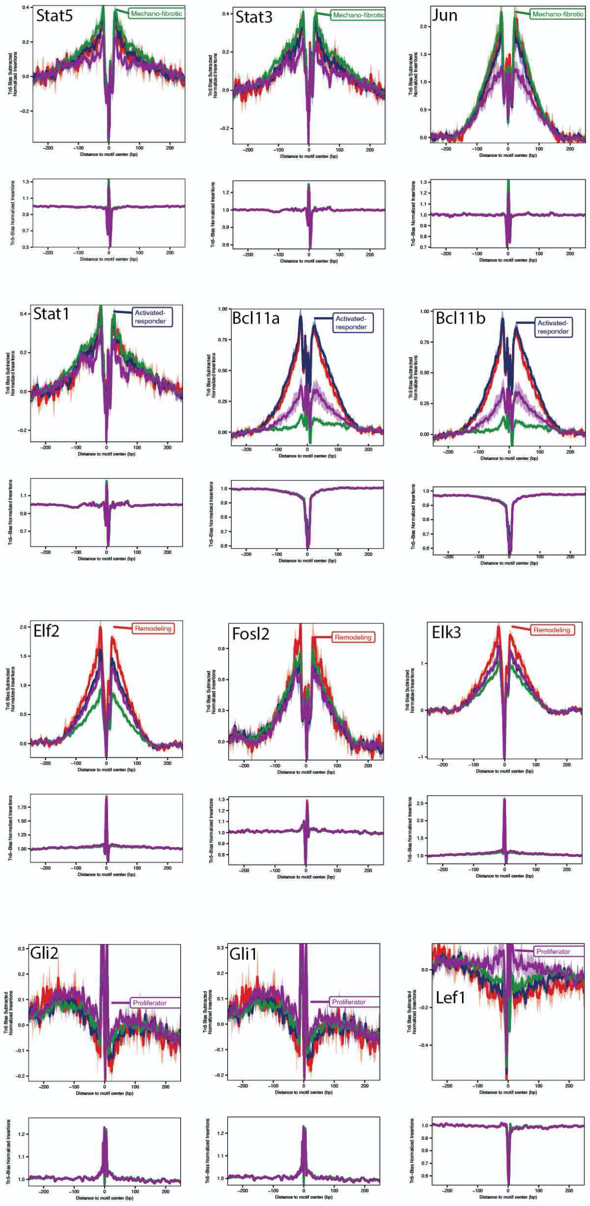

### Extended Data Figure 9

**a** GREAT (Genomics Regions Enrichment of Annotations Tool) Analysis

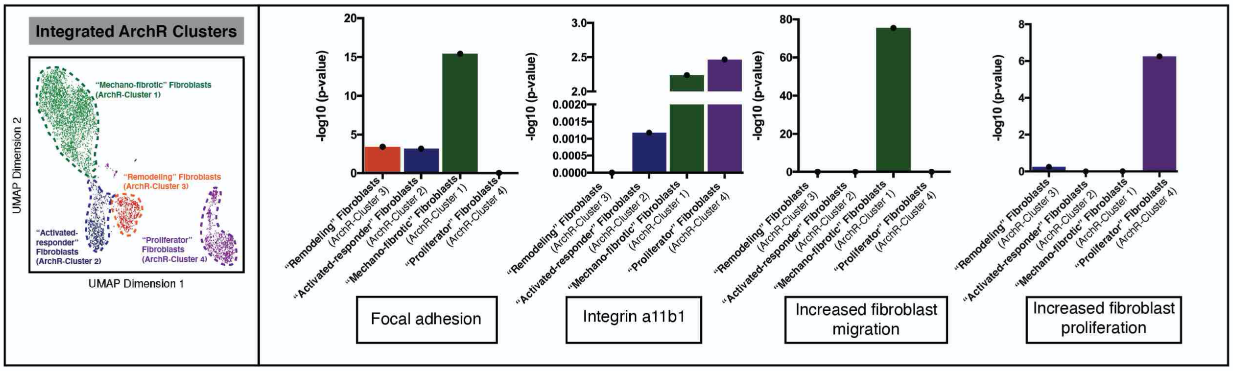


**b**

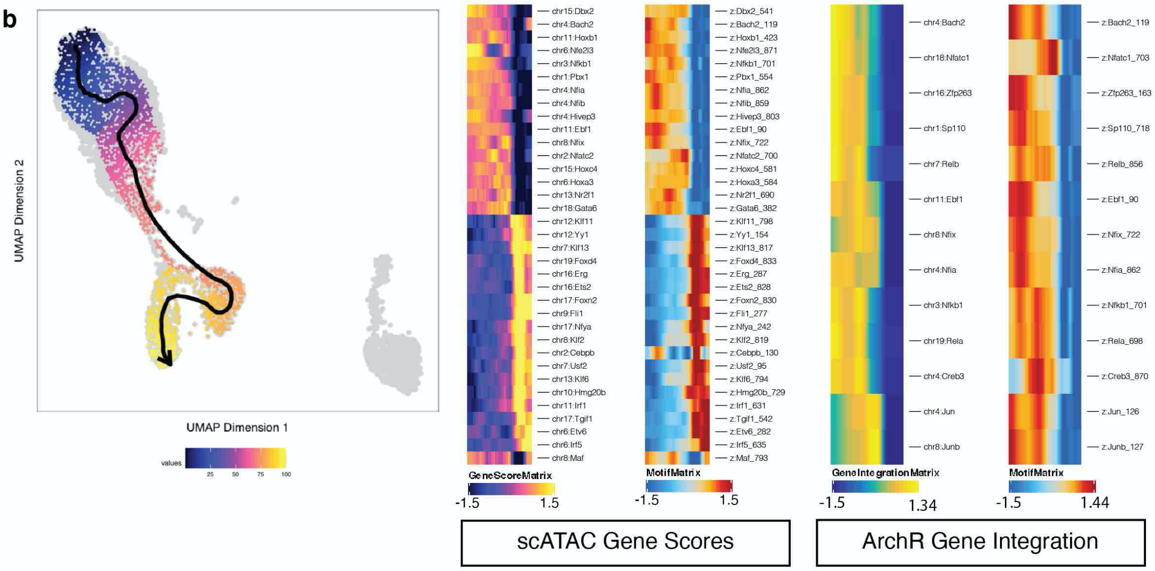

### Extended Data Figure 10

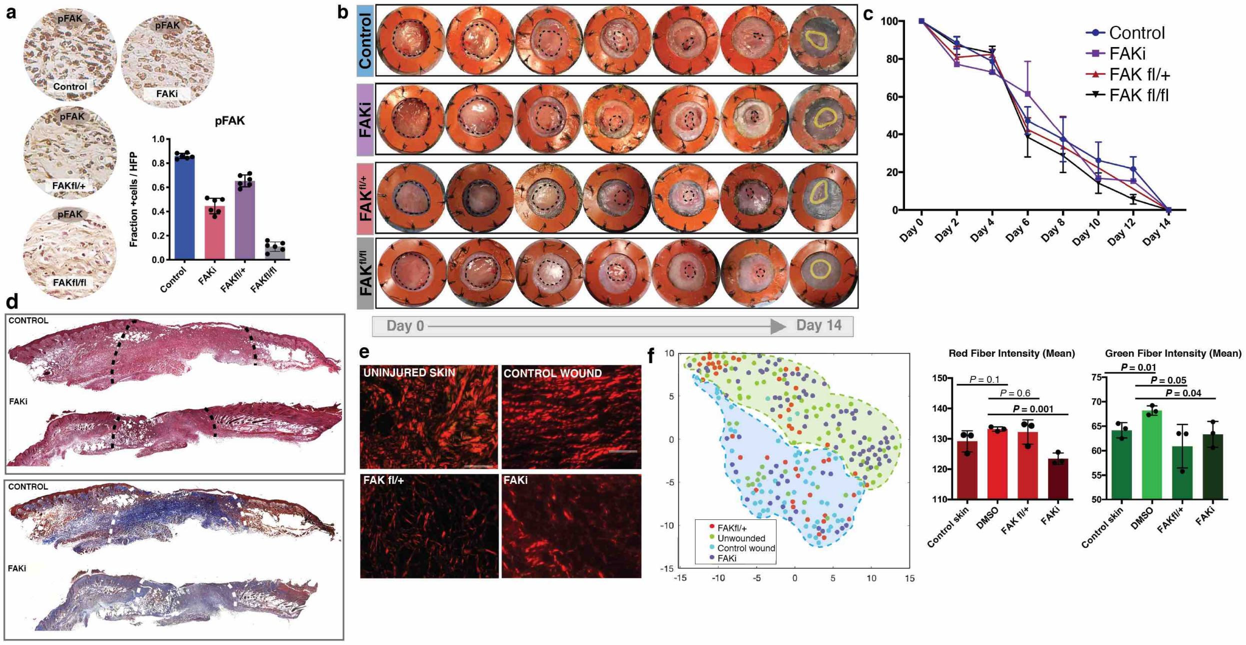

### Extended Data Figure 11

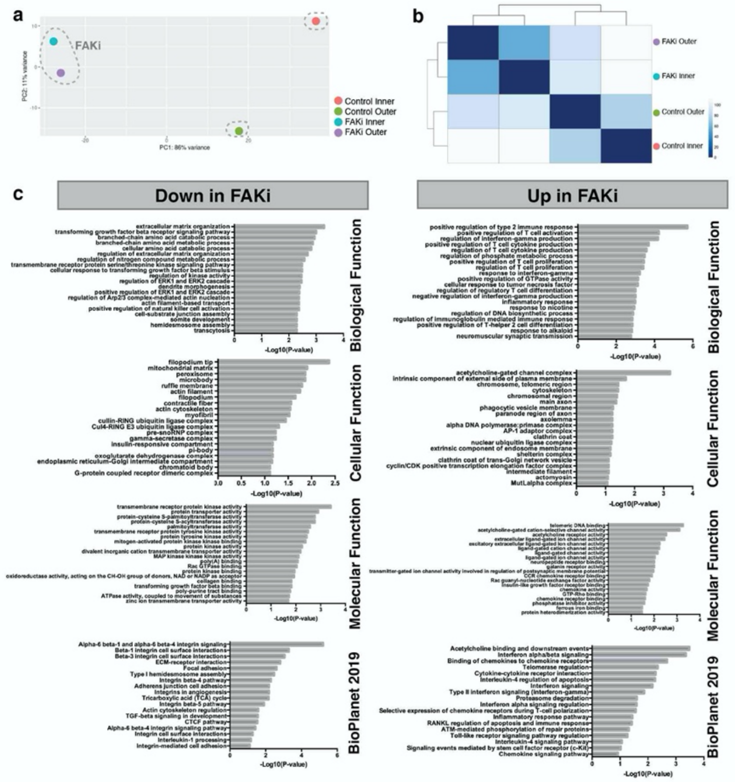

**Extended Data Figure 12**

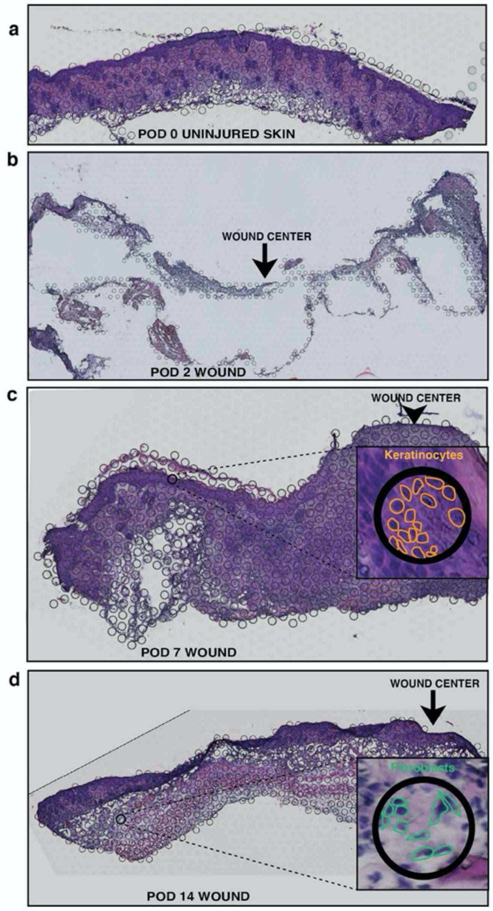

Extended Data Figure 13

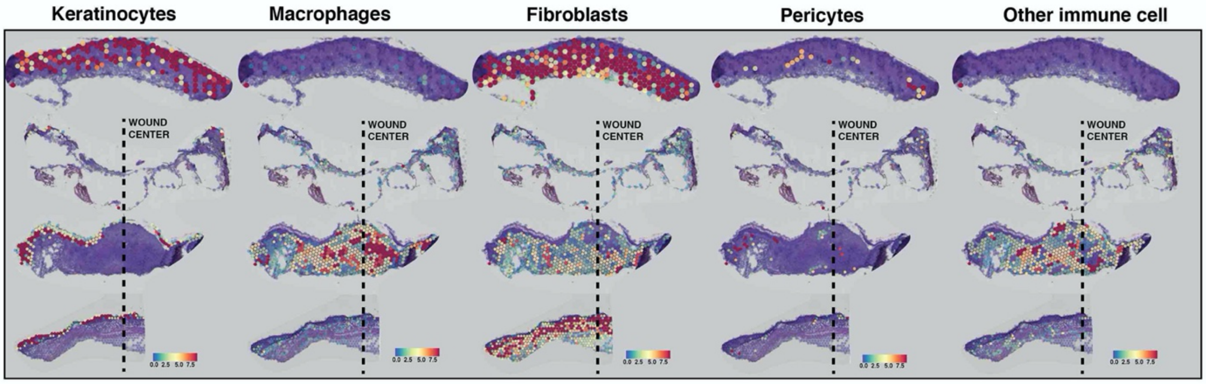

**Extended Data Figure 14**

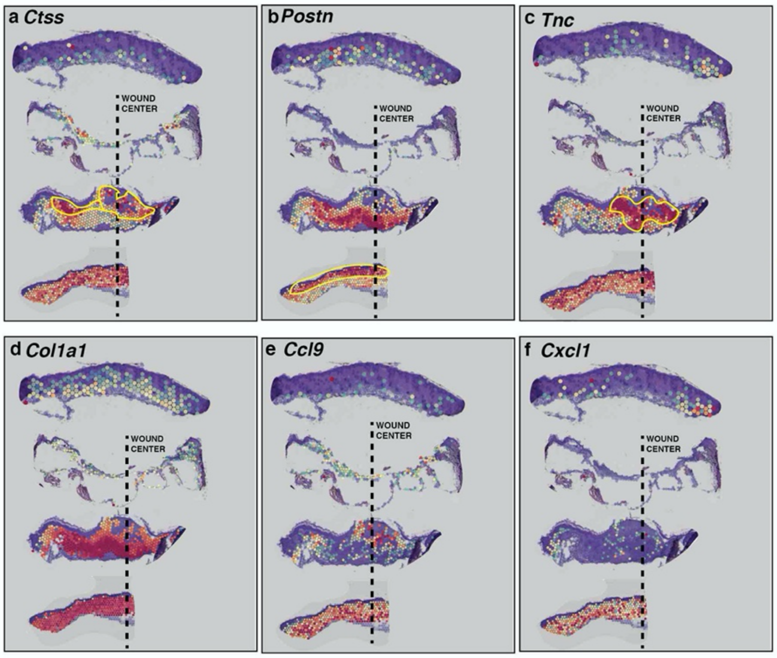

**Extended Data Figure 15**

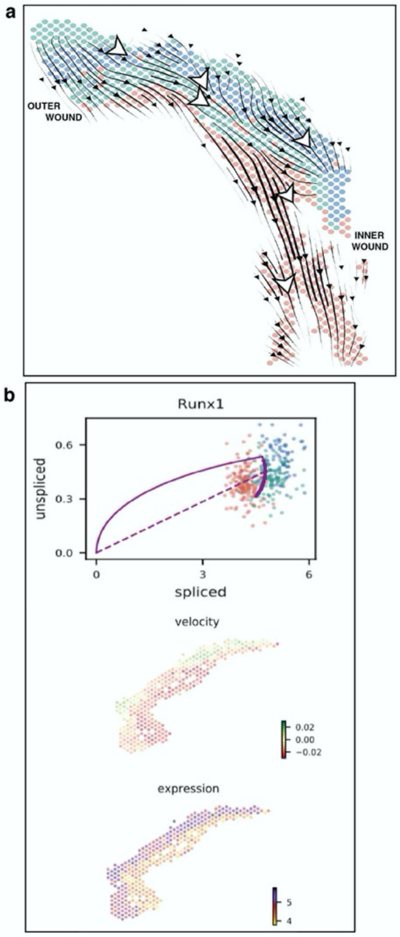

**Extended Data Fig 16**

a

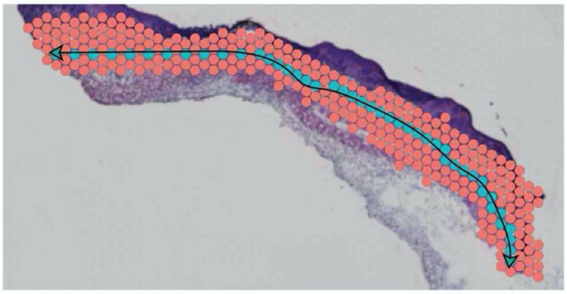

b

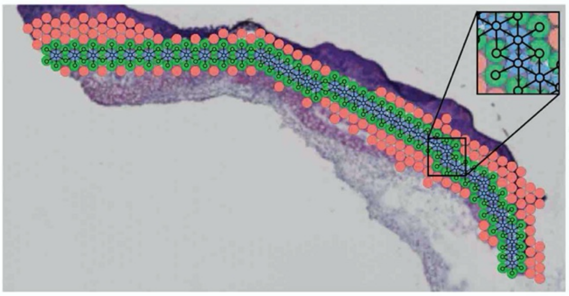

Extended Data Figure 17

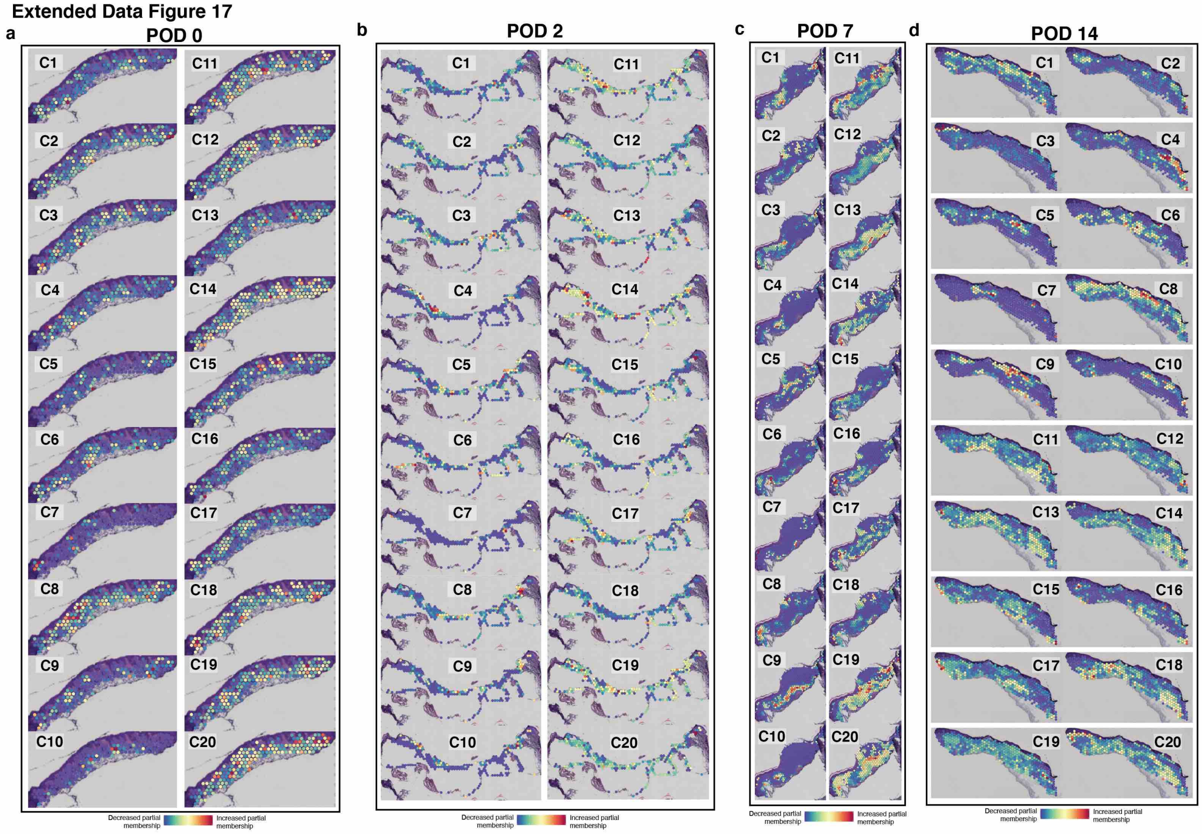

Extended Data Figure 18

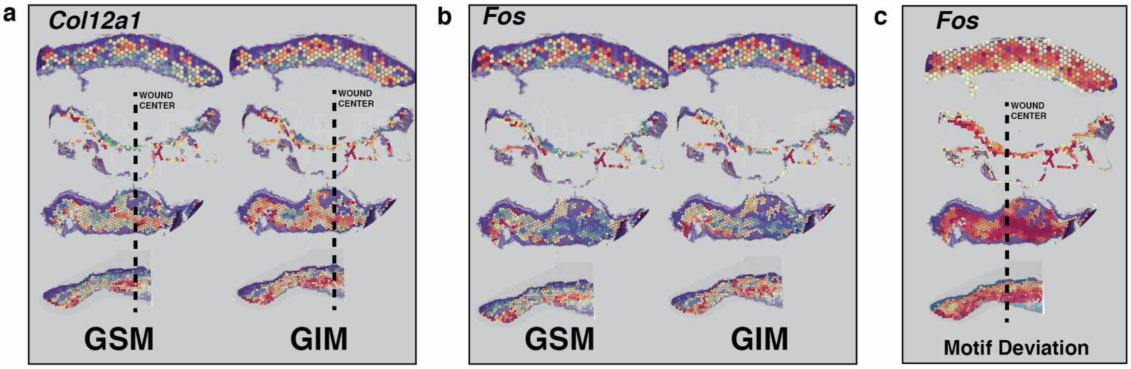

### Extended Data Figure 19

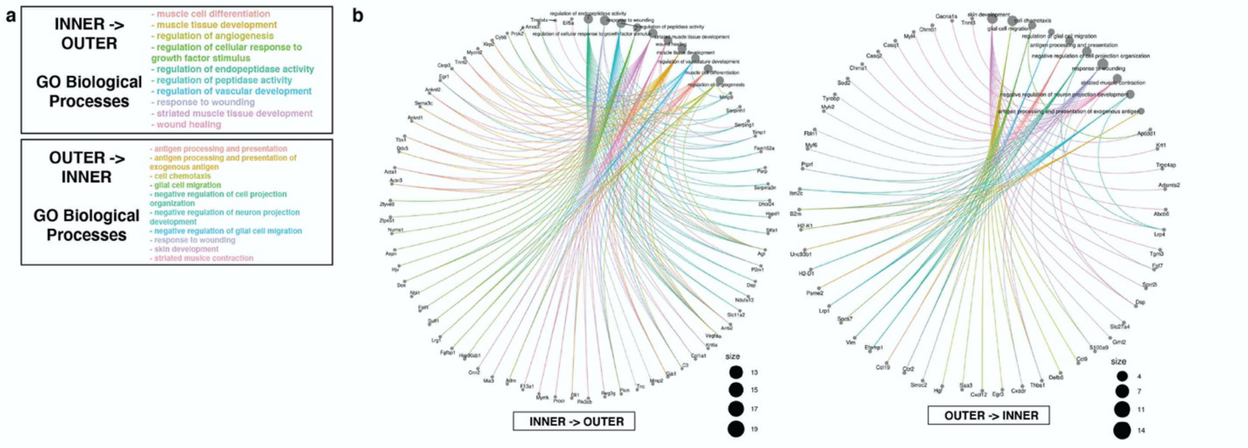

**Extended Data Figure 20**

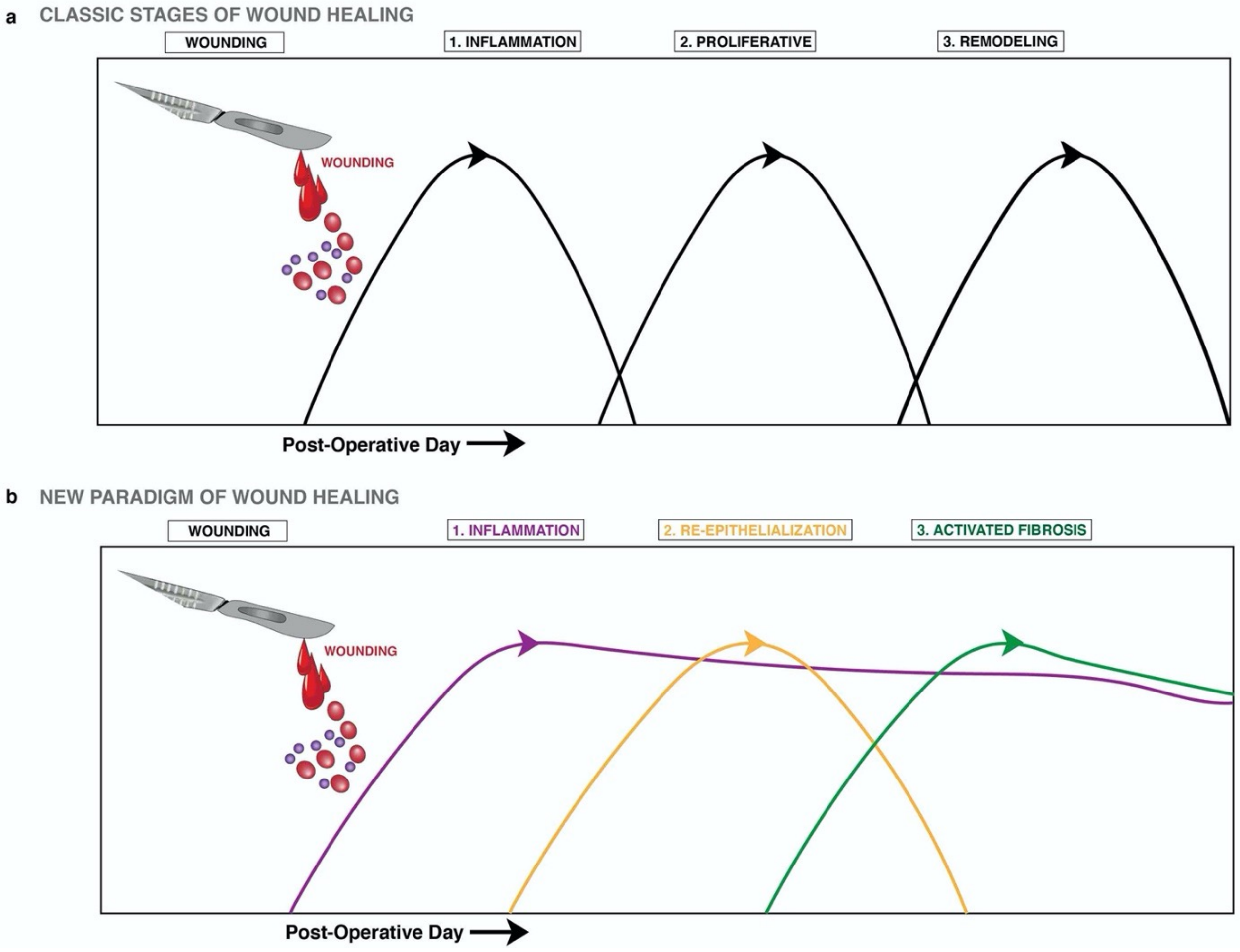
